# Molecular Dynamics Reveal Base Flipping as a Key Mechanism in Tc3a Transposase DNA Recognition

**DOI:** 10.1101/2025.03.27.645863

**Authors:** Stephan L. Watkins, Helder I. Nakaya

## Abstract

Transposons, or “jumping genes,” are mobile genetic elements that reshape genomes. The Tc1/Mariner family, including Tc3a transposase, has been studied for decades, but key aspects of its DNA recognition and cleavage remain unresolved. Existing models propose a homo-dimeric process for DNA excision and integration, yet conflicting hypotheses exist about the initial steps of DNA cleavage and target site recognition. Here, we reveal a previously unrecognized base-flipping mechanism in Tc3a transposase that challenges long-standing models of Mariner transposon activity. Using molecular dynamics simulations, we demonstrate that N-terminal acetylation induces a structural shift in the transposase-DNA complex, forcing a nucleotide in the inverted repeat to flip outward. This base-flipping event alters the local DNA conformation, creating torsional strain that may facilitate transposon inverted terminal repeat association. Unlike prior models, which assumed the entire protein, or protein dimers, were needed to illicit DNA interactions necessary for any types of torsion, our findings indicate that transposase bipartite binding itself actively reshapes DNA structure. We further show that this mechanism is dependent on the bipartite helix-turn-helix domains, which differentially contribute to DNA stabilization and recognition. Our simulations also reveal that the processed N-terminus plays an unexpected role in modulating DNA binding affinity, contradicting previous assumptions that it was structurally inconsequential. Together, the two helix-turn-helix motifs act to propagate the force caused by base flipping along one direction of the cognate DNA double helix axis. The fact that previous structural studies lacked this amino acid modification may explain why this mechanism was overlooked. These results provide a new framework for understanding transposase-DNA interactions, highlighting how a single molecular modification can trigger major DNA rearrangements. This discovery not only redefines the Tc3a transposition process but also calls for a reassessment of long-standing Mariner family transposon models.

## Introduction

Transposons are small stretches of DNA capable of “jumping” to different regions of an organism’s genome, profoundly influencing genetic diversity and evolution (1,2). These mobile elements are now classified into various families, each characterized by inverted terminal repeats (ITRs) flanking a central region (1200–4000 bp) that encodes one or more transposase, the enzyme responsible for their mobility (3,4). Transposons have been identified in every organism studied, ranging from inactive pseudo-transposons to fully functional elements (3,5). Their ability to facilitate horizontal gene transfer across kingdoms—such as between Protozoa and Animalia, or even between Oomycetes and Bryophytes—underscores their evolutionary significance (6). Moreover, their simplicity has made them invaluable tools in genetics, with applications in model organisms like Caenorhabditis elegans, Drosophila melanogaster, and Danio rerio, as well as potential candidates for gene therapy (7–11).

This study focuses on the Tc1/Mariner superfamily of transposons, first identified in C. elegans in 1989 (6,12). The Mariner superfamily encompasses over 1000 members, predominantly found in worms and insects, although some have been integrated into the genomes of humans and other vertebrates (13,14). A hallmark of this superfamily is its simplicity, a single transposase mediates the entire transposition process, guided by the ITRs that flank each transposon. Structurally, these proteins share a conserved N-terminal helix-turn-helix (HTH) DNA recognition domain, a short linker chain followed by a second HTH and a catalytic C-terminal core. This catalytic core is conserved throughout Tc1/mariners, consisting of a DDE/D Mg²⁺ binding site responsible for the entire excision and insertion catalytic processes defining the transposon jumping (15,16). Intriguingly, the structural homology of Tc1/Mariner transposases to transcription factors in higher organisms and restriction enzymes in prokaryotes suggests a shared evolutionary history, likely influenced by horizontal gene transfer (7,17,18). In many cases, only the bipartite or smaller HTH motifs remain in the host genome, especially for actively transcribed protein descendants. In some cases the retention of only DNA binding portions with host proteins as fusion proteins, arising from mechanisms involved in the transposition process such as insertion within introns coupled with host splicing, occur. This indicates a separation of the roles for the different domains of this simple protein being conserved independently, leading to novel models of protein evolution incorporating horizontal gene transfer.

Transposition in Mariners is thought to occur via a homo-dimeric mechanism in which the transposase excises the transposon at the ITR ends and inserts it at TATA-like sequences elsewhere in the genome (19–22). While the insertion process is structurally defined, key questions about the initial DNA cleavage and target recognition remain unresolved (16,19,20,23). These include several differing models of how the initial DNA cleavage occurs, ranging from complex multimerization of 4 transposase into a large synaptic complex to single tranposase cleavage at ends, then dimerization after cleavage. Another undefined aspect is the localization of both ITR into the final structurally identified dimeric insertion complex. It is currently not known if this dimerized end product is the result of a complex cleavage process, and passive diffusion of both transposon ends, or if an active process is undergone to bring the ends together. This temporal aspects of the process has been studied by several laboratories with wet laboratory techniques, with discrepancies between research findings. It is not known if the ends are cleaved sequentially in a more complex process, simultaneously at either end, or if bringing both ends together is part of the enzymatic process itself. In non-mariner transposon processes a wide range of mechanisms exist, including base flipping, holliday junctions, and multimeric protein complexes which have been used in some cases as a basis for development of parallel theories for Mariners (24–27).

Here, we investigated the ITR recognition of the Tc3a transposase to provide new insights into DNA binding and specificity in the Mariner family. Despite high structural similarity, Tc1/Mariner proteins show divergent sequence homology, making them an excellent model for studying the balance between specificity and general DNA recognition (18,28,29). Our first aim was to uncover the kinetics and mechanisms underlying these interactions, shedding light on DNA recognition processes relevant to transcription factors and DNA-binding proteins in diverse organisms, including Homo sapiens. Further, we explored a novel DNA-binding modality that appears to be a foundational mechanism in these transposons, with implications for understanding DNA and RNA interactions across vertebrates, plants, and mammals. This interaction may be significant to a subset of Mariners based on N-terminal sequence homology, as well as many DNA binding proteins with similar N-terminus sequences. We characterize the identified novel DNA interactions in detail, with a focus on DNA structural changes and affinity. This interaction, mediated by proper N-terminal amino acid processing, induces DNA structural changes within the ITR that aid in the general understanding of Transposon excision reactions in Tc3a or related Mariners. Applied, this research also addresses a critical gap in engineering synthetic DNA-recognition proteins for biotechnological and therapeutic applications. Utilizing only 1-2 amino acid changes in the N-terminal HTH of the Tc3a, or even changing a few atoms, we show drastic effects in affinity can occur.

## Materials and Methods

### Starting Structural Models

Our initial structure was retrieved from the Protein Data Bank PDB id=1U78, and modified accordingly (30). Based on normal C. elegans and H. sapiens protein processing, the N terminal methionine was removed (31), and terminal proline was modified to include a terminal acetylation. From the PDB DNA, the bound DNA was modified to include the Tc3 ITR sequence, 5’-TGTGGGAAAGTTCTATAGGACCCCCCCG-3’ and complement, 3’-CACACCCTTTCAAGATATCCTGGGGGGG-5’, which encompasses the entire N-terminal protein recognition portion, using Pymol. Gromacs-GPU 5.1.15 was initially used, with an all-atom Amber ff99SB-ILDN force field. Our Resulting model was centered in a rectangular box with dimensions 100×56×180 Å, solvated with 31726 TIP3P water molecules, 1 Mg2+ ion (0.002 M), 91 Na+ ions (0.140 M), and 30 K+ ions (0.040 M), with counter Cl-ions for each positive ion to mimic a minimal cellular salt concentration. The resultant box gave a 13 Å solvent layer to the Z axis, 40 Å to the X and 60 Å in the Y dimension (32). This process was repeated for models having an N-terminal modification with cleaved methionine and no acetyl group (NH2 -Proline), the terminal three residues removed with an NH3 Gly terminal end, and a NH3 terminal methionine followed by alanine and glycine, and no acetylated processing.

### Molecular Dynamics Model Equilibration

The resulting models were energy minimized using steepest descent, and equilibrated to 310 Kelvin, 1 atmosphere by first generating velocities at 290 Celsius for 2 ns non-varient temperature and then pressure for 2 ns, then continuing runs with these trajectories as input for 5 ns raising the temperature to 310 Kelvin, with Nose-Hoover temperature coupling, and Parrinello-Rahman Pressure coupling for the solvent and DNA-Protein. Resulting structural models were then further equilibrated using unrestrained MD with three temperature, pressure coupling groups DNA, Protein and water plus ions, for 20 nanoseconds. For further experiments, this resulting simulation was allowed to run for an additional 50 nanoseconds to generate adequate sampling space for starting pulled simulations, and an additional 300 ns for general bound bipartite model analysis. Hydrogen bonding or structural comparisons were analyzed using Gromacs, APBS software and in the software VMD (33–36). Model comparisons were done using Gromacs, VMD and Pymol (37,38). For the Acetylated bipartite, the model was used to generate 3 initial starting points, end extended for an addition 300 ns to determine the lifetime of the flipped base alone.

### Free Energy Calculations

Gibbs free energy was calculated using WHAM according to published PMF protocols using the above models (39,40). Briefly, Umbrella sampling for 50 nanosecond extended equilibrated segments from unrestrained simulations were used to generate starting models with time points between 5 and 50 nanoseconds chosen using a random time point generator, for a total of 16, and 26 selected times. These extension were described in model equilibration, and each time point used for pull initial start points. This was repeated for a total of 4 times, for the 4 separate N-terminal models systems, with the starting time points selection unique for each N-terminus. For the processed acetylated bipartite only, first 16 time points were used to create a starting point with the first HTH comprising the first 46 amino acids, HTH1, as the applied pulled force in the Z direction, with the DNA restrained at the 3’-G53 and 3’-C28 terminal Oxygen atoms with 2000 kJ/mol in the x,y,z directions. A pull force was applied at 2000 kJ/mol nm^2^, pull rate of 1 nm per nanosecond to the HTH1 center of mass (COM), with a pull tolerance of 10^-06^ kJ/mol/ps, and the simulation was run with the settings V-rescale temperature coupling on solvent and protein-DNA, C-rescale Pressure coupling, long-range cutoffs set to 1.4 nanometers, neighbor list update set to 2.4 picoseconds, Particle Mesh Ewald long range electrostatics, using an all-atom Amber ff99SB-ILDN force field. Runs varied over 7.2-8 nanoseconds to reach the final unbound state. Bound hydrogen bonds were restrained using the SHAKE algorithm to constrain bond vibrations so we could use a time step of 2 fs (41). This protocol was repeated using amino acids 47-59 for the connecting loop domain as COM applied force, and again for the second HTH comprised of amino acids 60-104 , HTH2. This resulted in 16 pulled simulations for each domain. A final set of 26 pulled simulations was then conducted, again using randomized time point selection, with HTH1, the loop domain, and HTH2 all pulled from their COM simultaneously.

This final pulled simulation set up was repeated for each of the described models, using the same three pull groups, and in all frames saved at 1 ps intervals. All pulled simulations were done using Gromacs GPU version 2023.2 on 4 H100 GPUs and 48 core Intel 2.3 GHz servers on the Brazilian National Computing Laboratory (LNCC), or 16 cores, 1 H100 GPU on CENAPAD Brazilian super computers, using all-atom Amber ff99SB-ILDN force field, long-range cutoffs set to 1.4 nanometers, neighbor list update set to 2.4 picoseconds, with Particle Mesh Ewald long range electrostatics, V-rescale temperature coupling, and C-rescale Pressure coupling for GPU use (42). Restraints, coupling groups and run settings were the same for all pulled simulations. Data was analyzed in Gromacs using the Wham method with the three simultaneously pulled groups and individual pull groups, then used for energy plots (39,43,44).

### Further Analysis

Further analysis was done using Gromacs analysis tools, with Gromacs versions respective of the simulations. For Root mean square fluctuations (RMSF) with the bipartite, the 300 ps terminal portion of the pulled simulations was compared to 300 ps in the bound state before pulled simulation start at the respective Umbrella sampling window time point and used for figures. For the RMSF of the ITR DNA, the terminal 50 ns of the unrestrained equilibrated model were used. In both protein RMSF, the presented data is the mean average of all simulations. For DNA and Protein, these RMSF were then changed from a minimum and maximum of 0 to 10 Å, and represented as 0-100% as B-factors for figures using gromacs.

For dynamic hydrogen bonding and density plots the mean average was calculated from the last 50 ns of data across the equilibration run, excluding the equilibration temperature and pressure normalization (45,46). Density plots represent averages with the maximum set to 18 nm^3^ , where all base pairs had a maximum between 17 and 19, with the exception of base pair 7-49 which had a maximum of 20. All density plots are scaled from 0.1nm^3^ at 0% to 18nm^3^ representing 100%, with pixel resolution at 0.001 x 0.001 nm. Density is measured in the x-y plane, with z spacing of +/-3.5 Å to incorporate all movement, and the center axis defined as the point between the phosphate groups of the pair. Centering of the projections in figures was done manually, from scaled output as a 5 nm x 5 nm window. The definition of nm^3^ is volume of highest occupied molecular orbital diameter of hydrogen per cubic nanometer, at the set temperature, which reduces to cubic volume, and represents motion of the base pairs respective of their defined center axis in the x-y plane over time respective of their atoms volume (45,47,48).

The software APBS was used to calculate the solvent accessible surface charge for bases T8-A16, C41-A48 and protein residues acetyl-Pro2-Arg3, and Arg54, using 150 mM ionic/water as dielectric solvent, at 310 kelvin. Images were scaled using eV, which are the expressed unit of K_b_T/e, the temperature corrected Boltzmann charge, and represented as charge shaded solvent accessible surface for DNA. Amino acids are represented as stick models for clarity.

For transition hydrogen bonding and distance, averages were produced from the entire pulled simulation sets, with only the 3 simultaneous COM pulled simulation sets used. A representative distance plot was used in figures as averaged plots vary significantly in starting time points or duration, causing the data to be lost in noise from averaged plotting of means. For average, minimum and maximum minor and major groove changes individual phosphate distance were calculated using 50 ns of the extended 300 ns unrestrained model, and groove dimensions calculated using the well described formulas outlined for DNA groove dimension calculations (49–51). Model snapshots, and trajectories were further analyzed, and graphic representations or figures made using VMD, Pymol, Qtiplot, Gimp and Weblogo (37,52).

### Protein Database Analysis

Protein Database Analysis was conducted as follows. The entire Uniprot protein database was downloaded as fasta files, and separated into Swiss-protein, and Tremble data sets. These were parsed with python scripts to tabulate and bin the fasta files containing N-terminal Pro-Arg. Further analysis was done using the resulting fasta files to assign percentages, classified according to organisms as listed in the Uniprot database, and generate tabulated data using Pfam, Interpro and GO based protein functional identifiers to assign protein function (53–56). Statistics were done based on direct DNA or RNA binding protein identifiers present in either database. Using only H.sapiens Uniprot entries, data was again parsed using several variant N-terminus chosen randomly or semi-randomly using a random number generator and the numbers 1-20 to represent amino acids for 5 N-terminus. Two of these were chosen intentionally to contain proline at position 1. Parsed H.sapiens data using the N-terminal Pro-Arg, Pro-His, Pro-Gln, Ala-Leu, Ser-Arg, and Arg-Gln to extract all fastas, was reduced based on presence of descriptive functional annotation and to remove redundant transcript variants, and enrichment analysis preformed with DAVID and MOET (57,58). The randomly generated terminus Met-Arg-Gln is the only one not processed into a terminal acetyl.

## Results

Initial equilibrated model structure hydrogen bonding varies from the X-ray structure, mostly through expected thermal expansion from the cryogenic determined atomic positions. With X-ray or EM models data is usually collected at -180 Celsius, leading to a 3-4 angstrom radial expansion for globular proteins of 300 amino acids, from center of mass. Our starting model and simulation setup are shown in Fig. 1. Main differences were Tyr46 made no hydrogen bonds with the DNA in any simulations, along with the Pro2 and Ala6 backbone nitrogens found in the X-ray structure. Our initial model also showed slightly different hydrogen bonding between the two helix-turn-helix (HTH) domains, referred to as HTH1 for the N-terminal and HTH2 for the C-terminus, as well as the connecting loop analyzed over 50 ns, from a 300 ns simulation. This loop sits more deeply in the widened minor groove, and makes different hydrogen bonds due to dynamics not completely included in the static structure mainly from arginines and lysines (30,59). However the bulk of difference was from the loop backbone, where in the dynamic simulations the nitrogens of the backbone chain hydrogen bonds with the DNA phosphate backbone between residues 50, 51, 52, 55, 57, and 59, as well as a Pro56 N to C41-H1’ bond. Residues are number starting with the cleaved N-terminus methionine or acetyl group as 1 in the work presented here, which correspond to the same numbering in published X-ray work. A complete list of hydrogen bond differences is presented in S1 Table, all atom names used follow the AMBER all atom force field nomenclature.

**Figure 1:**
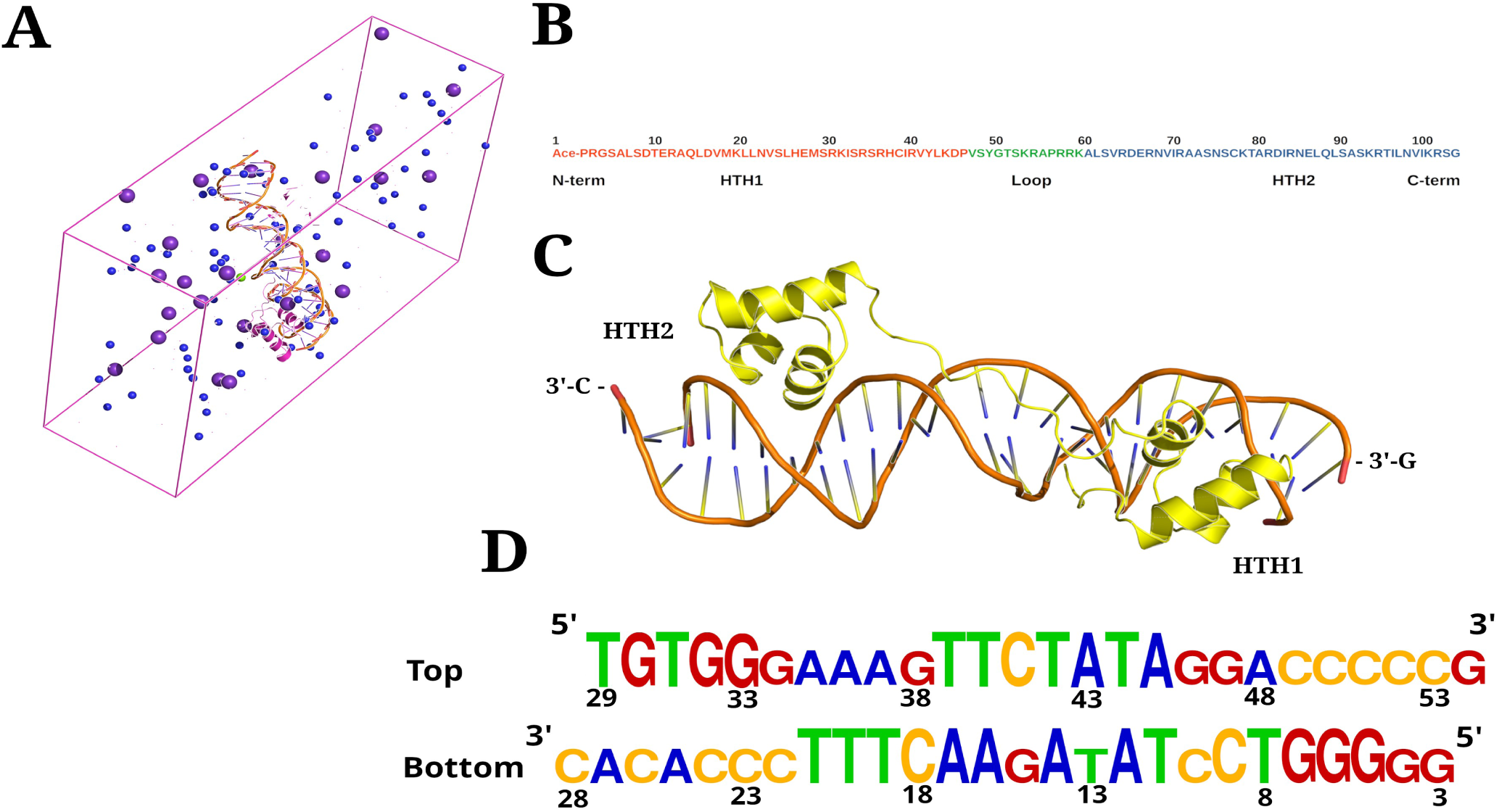
Initial Pulled Simulation Model. A) A representation of the unit cell, with the bipartite centered in the middle, ions K, purple, Na, blue, Mg green, Cl and H2O is not shown. The force was applied along the long axis, and simulations run until half the protein entered the periodic wall. B) Amino acid sequence of the bipartite, red HTH1, green loop, blue HTH2 C) Cartoon representation of the equilibrated starting model, N-terminus and HTH1 is to the right. D) The sequence of the dsDNA, aligned to the model in C), larger letters indicate a hydrogen bond contact with the bipartite protein.

For the dynamic process, beginning with the acetylated-Pro, the residue is almost stationary in position with respect to the ITR DNA, however moves lightly more than all other N-terminus tested. The Acetyl oxygen forms a optimized repulsive interaction at 3.4 Å with the ring N3 of A12, forming a perfect 109° angle with the sugar O4’ of T13, and O4’ to N1 of A43 angle at 121.6° and 4.0 Å. A slight 3.3 Å hydrogen bonding to the almost neutral H2 of A43 may help stabilize this reaction, although the bonding only occurs transiently 6% of occupied time, or with a lifetime of only 2.4 ps, along with 3% and 3% respectively for H4’ and H5’1 of T13 sugar ring, each with a 1-1.2 ps lifetime or less. This position is more reinforced by the proline’s almost complete hydrophobicity due to the neutralizing nitrogen modification, pushing the residue into the center of the DNA, and being completely solvent exposed on one side. This hydrophobic effect also includes the CH3 of the acetyl group itself. Residue Arg3 forms an antiparallel base pair through NH2 hydrogen and the sugar of T11 O4’, trapping the base O2 which is also bound by the hydrogen of the NE. This places the Arg main-chain N within 2.5 Å of the complementary base, A45, where base N3, formed a bond occupied 68.3% of the time, and a salt bridge with O1 of T13 phosphate, mediated by a fixed H2O, 50% occupied through simulations.

Together this forces a contraction of the minor groove on one side towards HTH1, and a widening, aided by Arg54 immediately below the modified Pro2 in the adjacent direction. These bonds, Arg54 NH2 hydrogen with A43 base N3, Arg54 NE hydrogen with A43 ribose O4’, and Arg54 NE hydrogen with T42 base O2 are occupied extensively in simulations. This interaction pulls slightly on and stabilizes A43, opposite T13. Together, the acetylated N-terminal Pro-Arg3 and Arg54 cause the T13 to adopt a flipped out conformation rotating 120-140 degrees into the solvent. This conformation, not observed in X-ray data, is maintained through the duration of separate long simulations for 300 ns. The charge distributions, and structure of the region are shown in Fig. 2, highlighting the effective use of hydrophobic force and stabilizing hydrogen bonding to cause base flipping in small HTH containing DNA binding proteins. All hydrogen bonds lasting longer than 1 ns, and having an occupancy higher than 40% for longer bound simulations are presented in Table 1.

**Figure 2:**
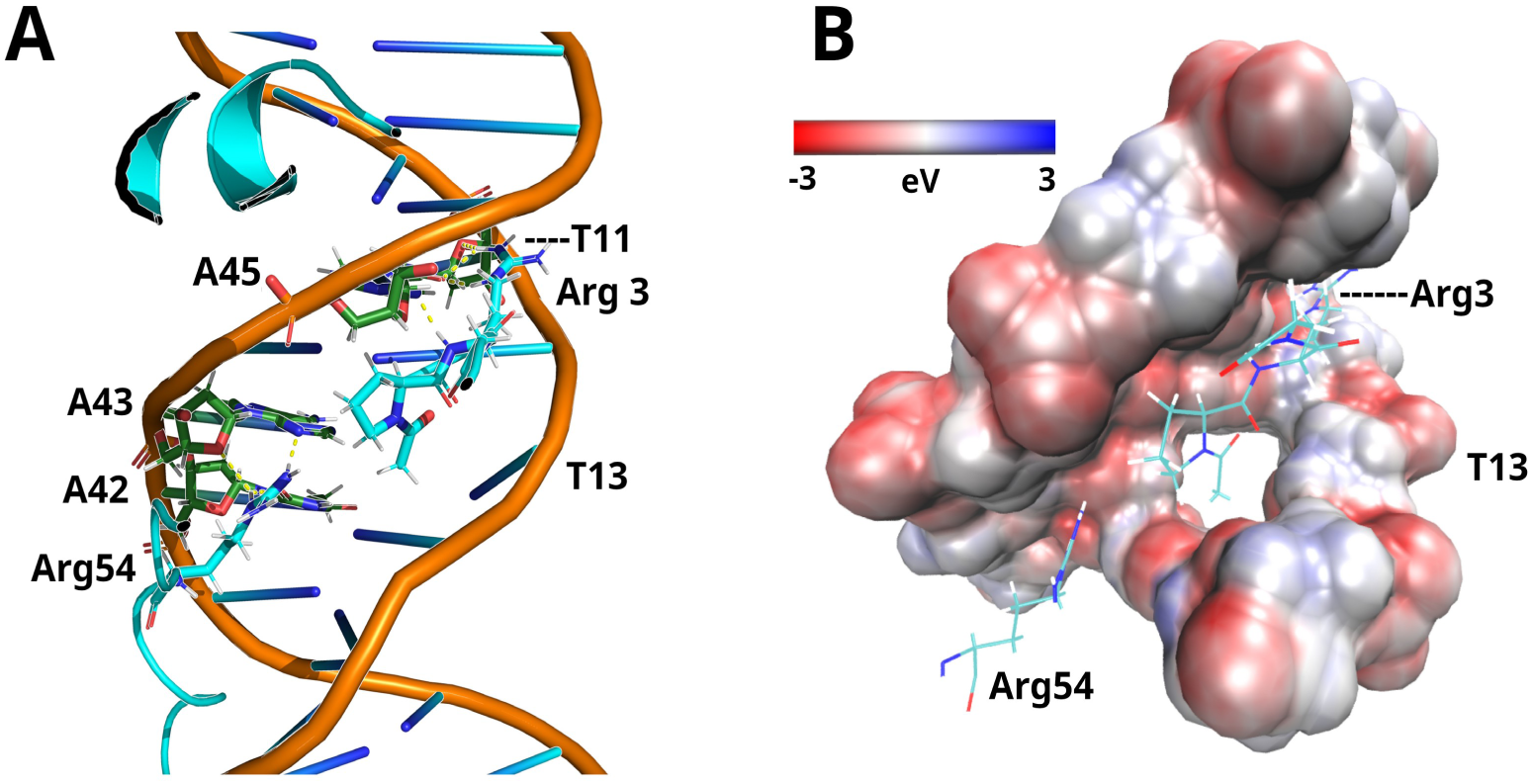
N-terminus Binding Site. A) Cartoon of the binding site for the N-terminus, including Arg54. Bases are labeled with 1 letter DNA code and position number. Hydrogen bonds are colored as yellow dotted lines. B) Same orientation as A), DNA shown as space filling model with APBS calculated partial charges shown according to scale as red, negative and blue, positive. Protein is shown as stick model, colored by atom type, red, oxygen, blue nitrogen, carbon light blue, hydrogen white. A hole formed through the DNA from flipped T13 is occupied by the Acetyl group.

**Table 1.** Long Lived High Occupancy Protein-DNA Hydrogen Bonds

| Functional Group |  | Percent Occupancy |
| --- | --- | --- |
| H-Donor | H-Acceptor |  |
| ARG3-NE | T11-O2 | 99.00% |
| ARG3-NH2 | T11-O2 | 78.34% |
| ARG3-NH2 | T11-O4' | 7.85% |
| ARG3-NH1 | A12-O4' | 21.06% |
| SER24-OG | G5-O1P | 94.95% |
| SER24-OG | G6-O1P | 11.65% |
| HIS26-NE2 | G6-N7 | 80.85% |
| HIS26-NE2 | G5-N7 | 6.07% |
| ARG30-NH1 | G5-O1P | 99.03% |
| ARG30-NH2 | G5-O2P | 98.53% |
| ARG34-NH1 | G46-O1P | 96.85% |
| SER35-OG | G46-O2P | 99.10% |
| ARG36-NH1 | G7-N7 | 99.70% |
| ARG36-NH2 | G7-O6 | 93.87% |
| C9-N4 | HIS37-ND1 | 52.39% |
| HIS37-NE2 | G46-N7 | 22.52% |
| HIS37-NE2 | A45-N7 | 6.92% |
| CYS38-HG | A45-O5' | 10.62% |
| CYS38-HG | G46-O2P | 7.87% |
| ARG40-NH2 | G7-O2P | 81.53% |
| ARG40-NH1 | T8-O2P | 17.39% |
| SER52-OG | A44-O2P | 44.89% |
| SER52-OG | A43-O3' | 12.85% |
| LYS53-NZ | A16-O1P | 5.45% |
| LYS53-NZ | A17-O1P | 35.32% |
| ARG54-NH2 | A43-N3 | 99.10% |
| ARG54-NH1 | A43-O4' | 11.29% |
| ARG54-NH1 | A14-N3 | 7.6% |
| ARG54-NE | T42-O2 | 48.91% |
| ARG57-NE | T39-O2 | 32.69% |
| ARG57-NH1 | C18-O2 | 26.77% |
| ARG57-NH2 | T40-O1P | 8.57%% |
| ARG58-NH2 | T42-O2P | 45.24% |
| ARG58-NH1 | T41-O2P | 35.44% |
| ARG58-NH2 | C41-O3' | 47.31% |
| LYS59-NZ | C18-O1P | 21.67% |
| THR79-OG1 | T31-O2P | 76.06% |
| ARG81-NH1 | G30-O2P | 52.01% |
| ARG81-NH2 | T31-O2P | 48.69% |
| SER92-OG | T20-O2P | 47.24% |
| SER92-OG | T20-O5' | 47.22% |
| LYS93-NZ | G32-O6 | 60.98% |
| LYS93-NZ | G33-O6 | 39.11% |
| ARG94-NH2 | T21-O4 | 9.42% |
| ARG94-NH2 | G34-N7 | 54.29% |
| THR95-OG1 | T20-O2P | 97.15% |
| LYS101-NZ | G33-O2P | 36.19% |
| ARG102-NH1 | G33-O2P | 98.31% |
| ARG102-NH2 | G33-O2P | 98.27% |
| **Backbone** |  |  |
| ARG3-N | A45-N3 | 68.01% |
| ARG3-N | T44-O4' | 9.80% |
| GLY4-N | A45-O4' | 14.15% |
| LEU25-N | G6-O1P | 76.63% |
| LEU25-N | G6-O2P | 4.47% |
| HIS26-N | G6-O2P | 59.66% |
| SER35-N | G46-O2P | 89.20% |
| GLY50-N | A45-O2P | 27.22% |
| GLY50-N | A44-O1P | 6.15% |
| THR51-N | A44-O1P | 33.82% |
| SER52-N | A44-O1P | 43.96% |
| ALA55-N | A43-O1P | 11.94% |
| C41-H1' | PRO56-O | 7.47% |
| ARG57-N | C41-O4' | 21.04% |
| ARG58-N | C41-O2P | 47.31% |
| LYS59-N | C18-O1P | 7.97% |
| ALA60-N | T19-O1P | 92.98% |
| ALA80-N | G32-O2P | 72.31% |
| ALA91-N | T20-O1P | 16.05% |
| SER92-N | T20-O1P | 48.42% |
| SER92-N | T20-O2P | 48.41% |

List of all hydrogen bonds between Protein and DNA with higher than 10 ps, 10% occupancy across extended simulation. Amino acids that meet the criteria once also have shorter bond occupancies listed. Functional Group, atomic interactions from amino side chain atoms, Backbone, atomic interactions from amino mainchain. In both column 1, donor, column 2 acceptor, column 3 percent occupancy across 100 ns pre-equilibrated simulation.

Distortions in the DNA only slightly mirror X-ray structures of Tc3a, Mos1 or related Mariners with a shift to A-DNA. The flipped T13 causes the phosphate of A14 to bend inwards towards the minor groove, however this torsion causes the minor groove between G15 to C17 and C41 to A45 to adopt a wider opening. Respective averages by base from G15 are 15.54 Å, with a maximum 19.34 Å, minimum 11.89 Å , to A17 with an average of 13.26 Å , and maximum 15.71 Å, minimum 11.32 Å. The highest distortion occurs from G15 and A16 paired with T44 and A45 on the opposite strand. Conversely, the opposite direction of the flipped T13 causes the minor groove between DNA residues from C9 to A12 and C51 to G48 to contract. Here the highest variation is on C10 and T11, with average, minimum and maximum of 10.26 Å , 6 Å, 15.20 Å and 12.33 Å, 6.56 Å, 15.51 Å respectively. Residues C9 and A12 have only minimums of 8.9 Å, with averages in normal B-DNA ranges. With both distortions, the widening in one direction and contraction the opposite stretch, the occurrences are sinusoidal over the simulations. Each holds positions outside of standard B-DNA conformations with the widening direction towards HTH2 having a lifetime of 250-300 ps, maximum 400 ps, and the contraction towards HTH1 a maximum life of 80 ps, and average of 40 ps. The widened minor groove between HTH1 and HTH2 is occupied by the Loop domain, and in contrast is contracted in the X-ray structure, whereas the TATA region in published X-ray structures is not comparable due to the base T13 flipping, the minor groove from T11-A14 has a 8.9 Å width in separate published X-ray structures. Additionally, this indicates the HTH2 rather than HTH1 covered regions are closer to an A-form of DNA through the simulations lifetimes. These highlight the further differences between static X-ray DNA-protein interactions and simulations at physiological conditions necessary for the study and development of DNA binding proteins.

Tc3a processed N-terminal amino acids have a strong effect on DNA affinity. We conducted pulled molecular dynamics to determine the binding affinity of the Tc3a bipartite domain, and the respective HTH1 and HTH2 individually, along with the connecting loop region. Total affinity was higher than expected from published work, where the average HTH has been shown to be around -80-120 kJoules/mol, although the total affinity of the bipartite was almost exactly the same for dozens of complete DNA binding transcription factors, around -400-440 kJoules/mol (60–62). Our simulations showed the entire bipartite of Tc3a Gibbs free energy change to be -416.17 k Joules, with the change in free energy presented in reverse of the binding process in Fig. 3A and S1A Fig. However analysis of the domains showed a normal, expected range for HTH2, -114.5 kJoules. Most of the energy could be attributed to the HTH1 domain and loop region, with a -155.7 and -148.2 kJoule/mol affinity respectively. This was higher for the HTH1 than expected, but was extremely high for a small 12 amino acid loop that in studied transposase only resides in minor grooves making primarily phosphate based interactions, Fig. 3D and S1D Fig.

**Figure 3:**
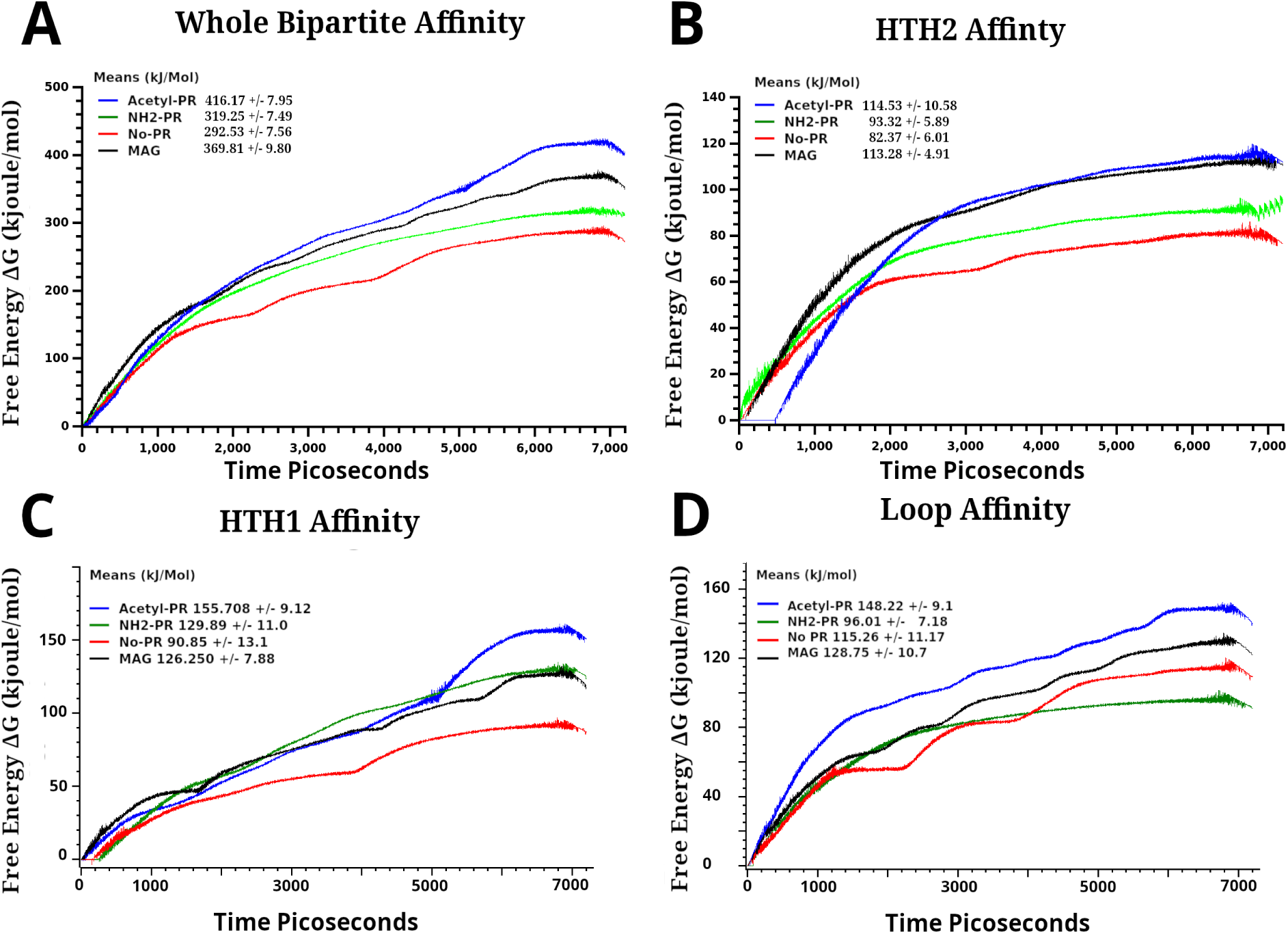
Free Energy Change. A) Binding affinity in kJoules/mol, and Gibbs free energy (ΔG) change of the entire bipartite domain averaged from all pulled simulations versus time, 7.2 ns, B) calculated mean affinity, and ΔG for HTH2, C) calculated mean affinity and ΔG for HTH1 , D) Binding affinity means for Loop region, residues Val47-Lys59, and ΔG over time. Final standard deviation are given as +/-either side of mean for A-D mean affinities in the legend. In all, Acetylated PR, blue, MAG, black, NH2-PR, green, and truncated to NH3-G end, red.

To further dissect the interactions responsible for the higher affinity, we repeated the pulled molecular dynamics simulations with changes to the N-terminal region. This resulted in 3 separate experiments, removal of the acetyl group replaced with NH2-Pro, removal of the terminal residues Pro-Arg with a truncated NH3, and replacement of the N-terminus with methionine, alanine and glycine as the terminal residues. Higher DNA affinity can be attributed to the acetyl group and primary 2 N-terminal amino acids. In all simulations, no large differences were found for HTH2 except with truncated removal of the N-terminus, as presented in Fig. 3B, S1B Fig. Here, the largest change for HTH2 affinity was from the removed Pro-Arg, with only Gly3-NH3 remaining. In these simulations, the entire bipartite is unbound by 5400-5600 ns, faster than all other tested altered bipartite, which will alone lower calculated free energy changes.

This is mirrored in our most profound change, found in HTH1,where removal of the N-terminal Pro-Arg resulted in a binding affinity of only -90.9 k Joules/mol. In contrast both the NH2-terminal-Pro and alternate Met-Ala terminus had roughly equal affinities at the higher range of expected HTH motif affinities from published research, while still lower than the acetylated N-terminus. Both the NH2-Pro and MAG had affinities of -129.9 and - 126.3 kJoules/mol respectively. Alone this showed the acetyl group itself contributes 25-30 kJoules to the HTH1 binding process, higher than can be attributed to any hydrogen bonding effects from a single oxygen, Fig. 3C, S1C Fig.

Unexpectedly, we also found extreme effects on the calculated loop affinities, Fig. 3D, S1D Fig. In contrast to the HTH1 the NH2-Pro structure had a much lower affinity, -96 kJoules, even lower than removal of the terminal residues. This was due to the NH2 and the proline preventing, or hindering one of the interaction of Arg54, placement of Lys53, due to alterations in backbone hydrogen bonding and backbone hydrogen bonding of the N-terminus. In simulations, the methionine N-terminus is able to partially occupy the same spaces as the natively processed N-terminus due to hydrophobic effects, without altering DNA structure, Fig. 4. This allows a Lys53 and Arg54 arrangement which was similar in the NH2-Pro terminal end simulations. The arrangement allows Arg54 to make hydrogen bonds with DNA bases A43 and G15, however the methionine methyl group prevents the Arg54 from entering deeper into the minor groove. In these simulations loop interactions are also hindered by the lack of slight widening of the groove found in the acetylated interaction. These Arg54 hydrogen bonds found in non-acetylated N-terminus models are rigid, with the methionine end having a constant hydrogen bond with both DNA strands, rather than a single side. This further prevents the loop from occupying space deeper in the groove, especially Ala55 and Pro56 which are driven by hydrophobic forces from the solvent. None of the other simulations flipped T13 out into the solvent, or any other bases in the ITR. Example visualizations of trajectory runs for each modified N-terminus and Acetyl-Pro-Arg are shown in S Video 1 to 4, with solvent removed.

**Figure 4.**
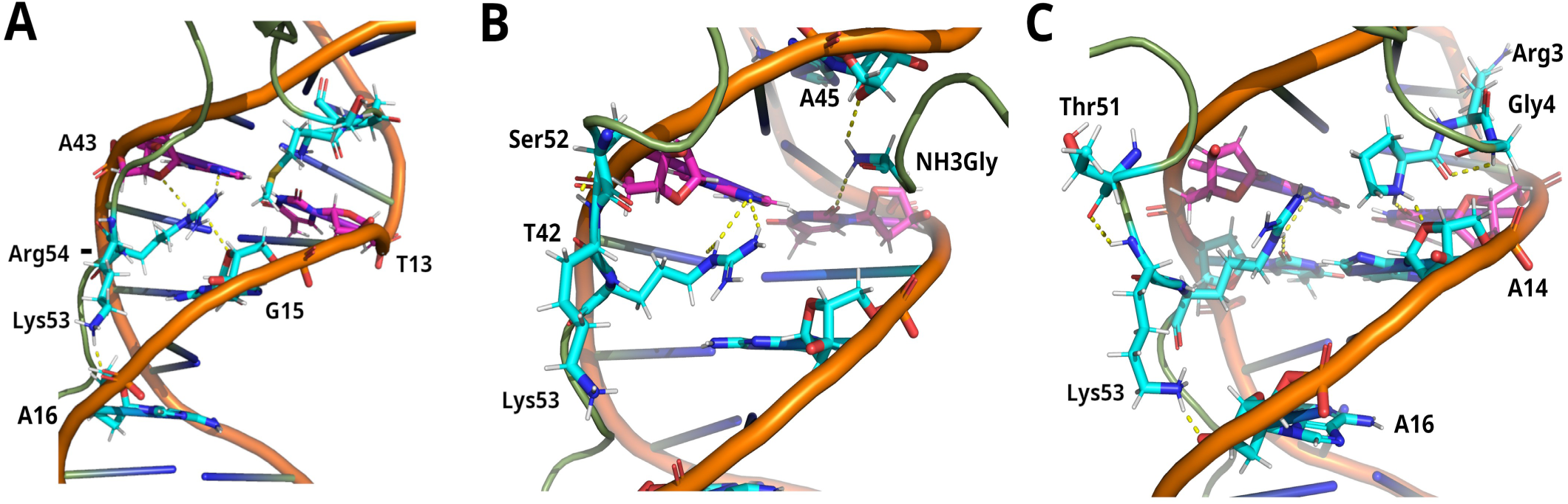
A)Methionine-Alanine-Glycine N-terminus Bound to ITR DNA. The modified N-terminus does not move deep into the minor-groove, and Arg54 has different hydrogen bonding than the unmodified terminus structure. B) Truncated NH3-Gly N-terminus hydrogen bonding and structure, C) NH2 Proline N-terminal hydrogen bonding and structure. All represent static structures highlighting long lived hydrogen bonding to illustrate main differences between the listed terminus, Lys53 and Arg54. Selected amino acids and bases are shown as stick models, while the remaining residues are shown as cartoon. T13 and A43 are fuchsia in all images, with carbons as fuchsia, while the remaining stick bases and amino acids are in aqua, with carbons aqua colored. In all, hydrogen, white, nitrogen, blue, oxygen, red. Hydrogen bonds are shown as yellow dashed lines. Lys53 branches above Arg54 towards the top of the image, but stretches downwards in all images. B,C are angled slightly different and slightly closer to the viewer due to differences in helix groove width, orientation of the amino acids and nucleotides, and to show hydrogen bonding. Each is centered the same as close as possible.

As a comparison, the truncated and NH2-Pro N-terminus show different hydrogen bonding effecting Lys53 and Arg 54, Fig4B and C. In the truncated end, the NH3 terminus of the glycine is held fixed by 3 hydrogen bonds, to T13 O2, A45 ribose O4’, and T44 O2. The Lys53 is unable to form hydrogen bonds with the DNA, mainly due to a kink formed by Lys53 NH to Ser52 O of the main chain. Also, Arg54 only forms bonds with one strand of the DNA, between a terminal amino NH and A43 ring N3, NH1 and A43 N3, with an addition bond between NH1 and T42 N3. This Arg54 hydrogen bonding is almost the same with the NH2-Pro N-terminal simulations, lacking the NH1 T42 N3 hydrogen bonding. However the kink shifts in this structure between the NH of Lys53 to Thr51 O main chain in the NH2-Pro structure, which also causes an almost helix like arrangement from Tyr49 to Lys53. This small change allows the lysine 53 to form hydrogen bonds with the NH3 end to A16 ribose O4’, and transiently with O5’ or O3’, much closer than the Acetyl and methionine structures. In the NH3-methionine structure Lys53 hydrogen bonds with only the O2-P of bases A16 and in the image, Fig. 4A, A17, rather than the ribose or base oxygens. Also the NH2 of the proline N-terminus forms weak bonds with T13 O2, and A14 ribose O4’, holding it more rigid than the Acetyl group. A small kink in the N-terminus is also formed by the Pro-main chain O and Gly4 NH2 not found in the acetylated or methionine structures.

To further analyze the effects on individual base pairs in the acetylated bipartite, we determined Root mean square fluctuations (RMSF) and density profiles for each base pair along the ITR over 50ns from 300ns simulations, Fig. 5, S2Fig. Both bp G4-C52 and C24-G32 show profiles close to what is expected geometrically in fluid simulations, with each base showing 80-90% of the space occupied by the ribose ring in a fixed position, Fig. 5B, S2D Fig (63,64). Base T13 remains in a flipped solvent exposed conformation the entire simulation, Fig. 5C, with much less movement in the position of the normally paired A43 base. Contraction of the minor groove in one direction causes a lengthening of the base interactions from bases A12 to C10, which is normalized around base C9 and T8. This repeats again for base pairs G5-G7, followed by normalizing at base G4. The majority of this lengthening for HTH1 is observed in bases T11 and A12, Fig. 5E, S2 Fig.

**Figure 5:**
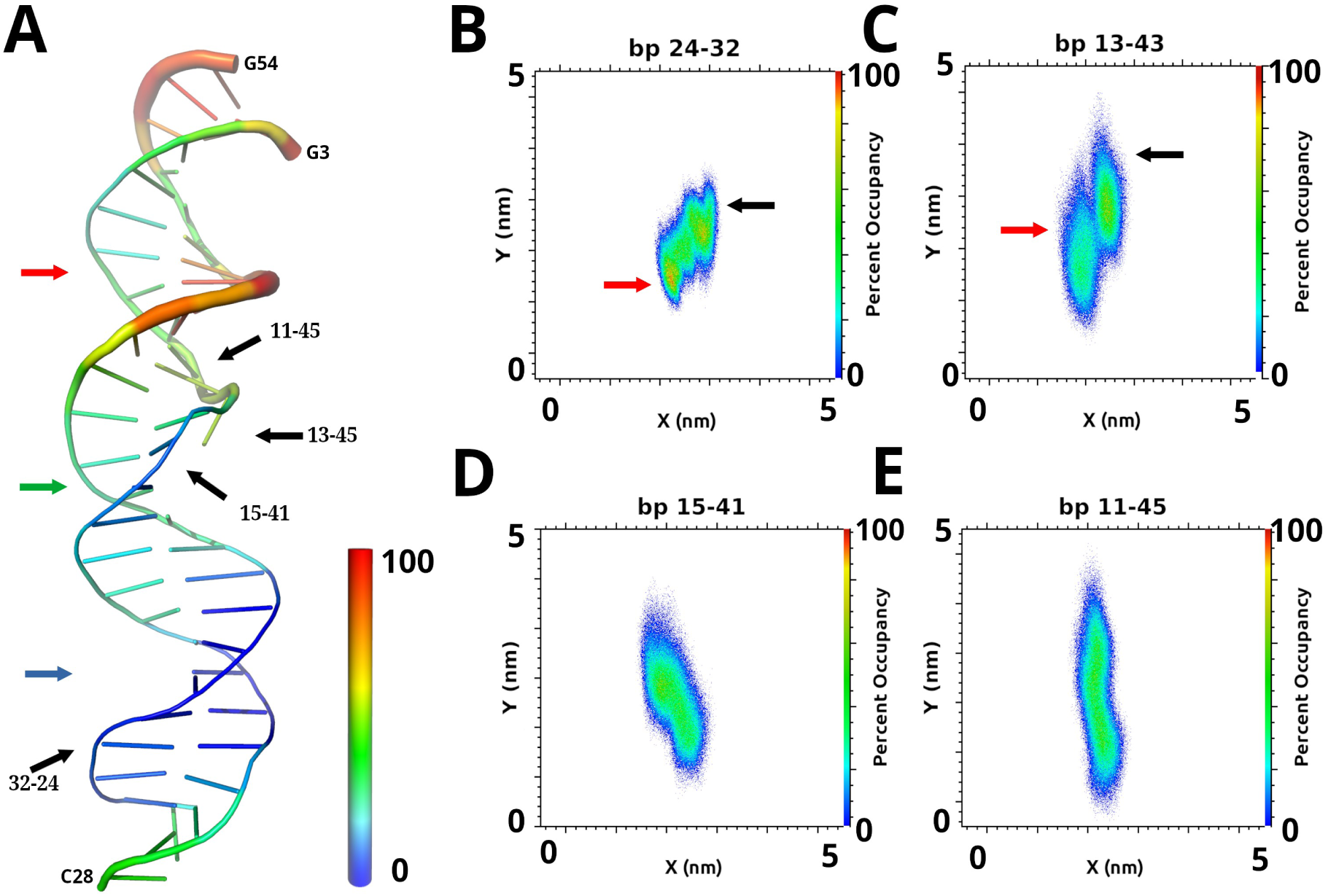
RMSF and Base Pair Densities for Extended Simulations. A) ITR DNA shown as jelly role, colored by motion normalized as B-factors from 0.1%-100%, thickness also indicates greater motion scaled to the same B-factor, representing normalized RMSF fluctuations over 50ns of extended non-pulled simulations. Red arrow, HTH1 binding site, green arrow, loop binding site, blue arrow, HTH2 binding site, specific base pairs from B-E, black arrows. B) Base pair C24-G32, shows an almost normal non-moving base pair density as reference, red arrow G32-P, black arrow C24-P C) the flipped T13-A43 base-pair, arrows indicate the positions of phosphates, black, A43, red, T13, D) G15-C41, showing a transition to less movement as in B), with slightly wider spacing, E) T11-A45, showing an extended spacing but still present centered base pairing of the terminal bases. In all, density a percentage of total space occupied for all trajectory frames in the x-y plane using the DNA central axis between base pair shown, including +/-3.5 Å in the Z plane to include the ribose and phosphate. Length across the long axis shows phosphate backbone positions, with the bases in between. Slight tilting of the bases also effects the width of density, and intensities, in the shorter axis.

In the opposite direction, the effect seems to mainly be with base pair interactions themselves moving slightly off at an angle or becoming extended, making the phosphate and ribose move more than the bases themselves from A16 to C18. This dissipates by base T19 to T20, but reappears more extreme from bases T21 to C23 with base C24 falling into normal expected DNA occupancy again. Base pair C26-G30 shows an extremely rigid profile not expected, S2D Fig, indicating it is almost fixed in position by HTH2, with the stretch C24-C26 showing extreme rigidity in the x-y plain. Together, movement profiles indicate base pairs from A16-C18 are moving more on one strand, namely paired bases T40-G38. Together, the RMSF and density plots determined show the overall x,y,z directions of movements induced by changes from the bipartite on the ITR. Almost no movement occurs in the Z direction taken as the length of the DNA towards HTH2, and large movements occur in the Z direction towards HTH1, which show a strong strand bias. This is also highlighted by the higher density for base pairs in the HTH1 region, indicating less x-y movement, whereas the HTH2 region seems to translate most motion into extended lengths between base pairs, truncated at a very rigid C26-G30 pair held by the bipartite.

Applying RMSF to the bipartite in a bound and unbound state revealed the minor dynamics of the protein domain. This showed results expected from simple HTH domain binding proteins, with HTH1 slightly more rigid in the bound conformation than HTH2, which contrasted the ITR fluctuations, Fig. 6. The Acetyl-Pro to residue Asp9 went from low to almost 90 percent fluctuation, meaning they are moving freely in the solvent even though forming an α-helix. Here the terminal 4 residues seem to rotate freely in solvent. This also translates into more springlike motion for the associated Helix one of HTH1, with the end Ser8 to Glu11 forming a helix that constantly extends then recoils. This is mirrored for the loop region from Tyr49 to Arg58, which is stabilized against the minor groove of the ITR through hydrogen bonding, with Lys53 showing the most movement. In simulations Lys53 locks over Arg54 to a phosphate oxygen only 40% of the time, further aiding the Arg maintain a one stranded hydrogen bonding conformation.

**Figure 6.**
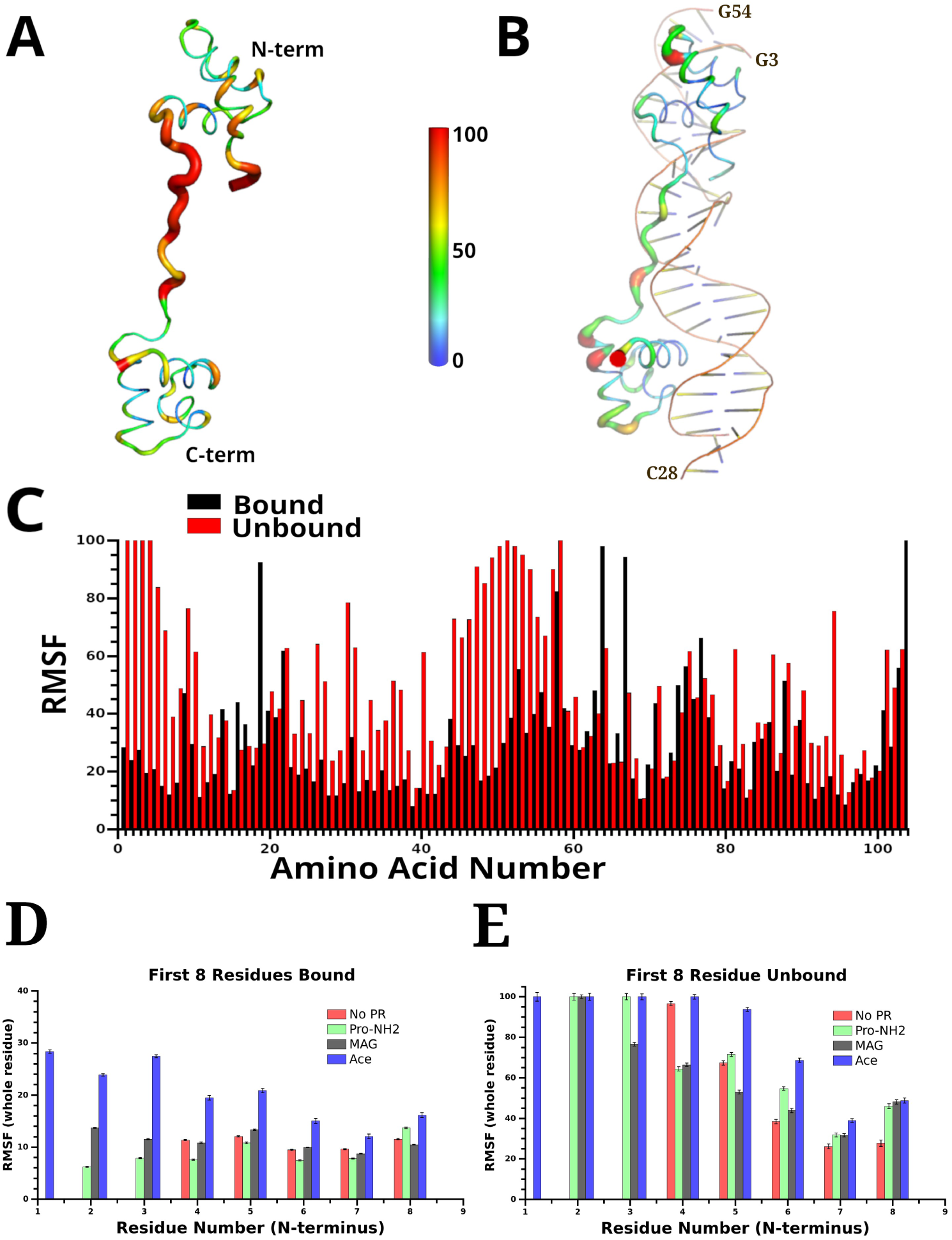
RMSF of Bipartite Domain. A) Jelly role representation of the unbound and B) DNA bound form of bipartite showing mean RMSF of all amino acids. B) shown as bound to ITR DNA, color, thickness represented as 0.1%-100%, 0% white, motion as normalized B-factor according to color scale shown. C) Normalized individual amino acids as bar graph scaled, as B-factors presented in A,B), bars in numbering at midpoint between bound, black and unbound, red. The last bar, residue 104. is the same for both conformations (red not shown). D) Comparison of the RMSF for bound and E) unbound 8 terminal residues from all 4 studied terminus. Residue 1, acetyl in C,D,E. For D,E) no PR is missing residues 1-3. Error bars represent STDV +/-. These are scaled as in C).

A remainder of motion changes are one of two, first the helices in both HTH become more compact, slightly rolling hydrophobic residues inward when unbound, allowing residues such as HTH1 Lys19 to Tyr42 and Lys31 to Thr10 or Asp9, to interact and hydrogen bond helix ends. This occurs with similarly placed residues on HTH2, where Arg64 and Arg67, are only solvent exposed in the bound form. Secondly, residues such as Arg30, Arg54, Arg40 or Arg 94 move from a strong DNA base interaction to being partially solvent exposed. These form less stable interactions with the adjacent amino acids in helices in the unbound form.

A comparison of the RMSF for all 4 N-terminus studied is presented in Fig. 6 D,E. Interestingly all other N-terminus had much lower RMSF in the bound or unbound form until residues 5 or 6 compared to the acetylated terminus. In the bound form, all 3 modified N-terminus behaved almost the same, with equivalent fluctuations 8-15% range. More divergence occurs in the unbound terminus, where the truncated NH3-Gly4 only shows higher fluctuation for the terminal 2 residues. For the NH2-proline, again only the first two residues show extreme fluctuation, while the MAG end only shows high fluctuation for the methionine, which also contains an NH3 end. Overall the MAG shows a more compact helix, starting from the alanine, while by residue 6 the truncated helical end, starting with NH3-Gly4 shows lower fluctuation in the helix overall compared to all other N-terminus.

To determine the DNA base pair interactions giving rise to or affecting the higher affinity, we looked at hydrogen bonding across all pulled molecular dynamics simulations, averaged across all runs. Initially we looked for only the Acetyl-Pro-Arg N-terminal interactions, and extended this to include Gly4 for hydrogen bonding, Fig. 7. Between 3800-4800 ps, the hydrogen bonding becomes solely Arg3, Gly4 peptide chain N and Ser5, and ends with a 2-400 ps stretch where only the Arg3 and Gly4 peptide chain N make bonds longer than 1 ps. A number of transient bonds are made from the Arg3 functional NH1, NH2 and NE hydrogens with T45, T13, T44 including the base oxygens or nitrogens, ribose O4’, and O1P or O2P lasting 1 ps or less. These interactions however are unable to account for the energy involved based on hydrogen bonding alone (43,65).

**Figure 7.**
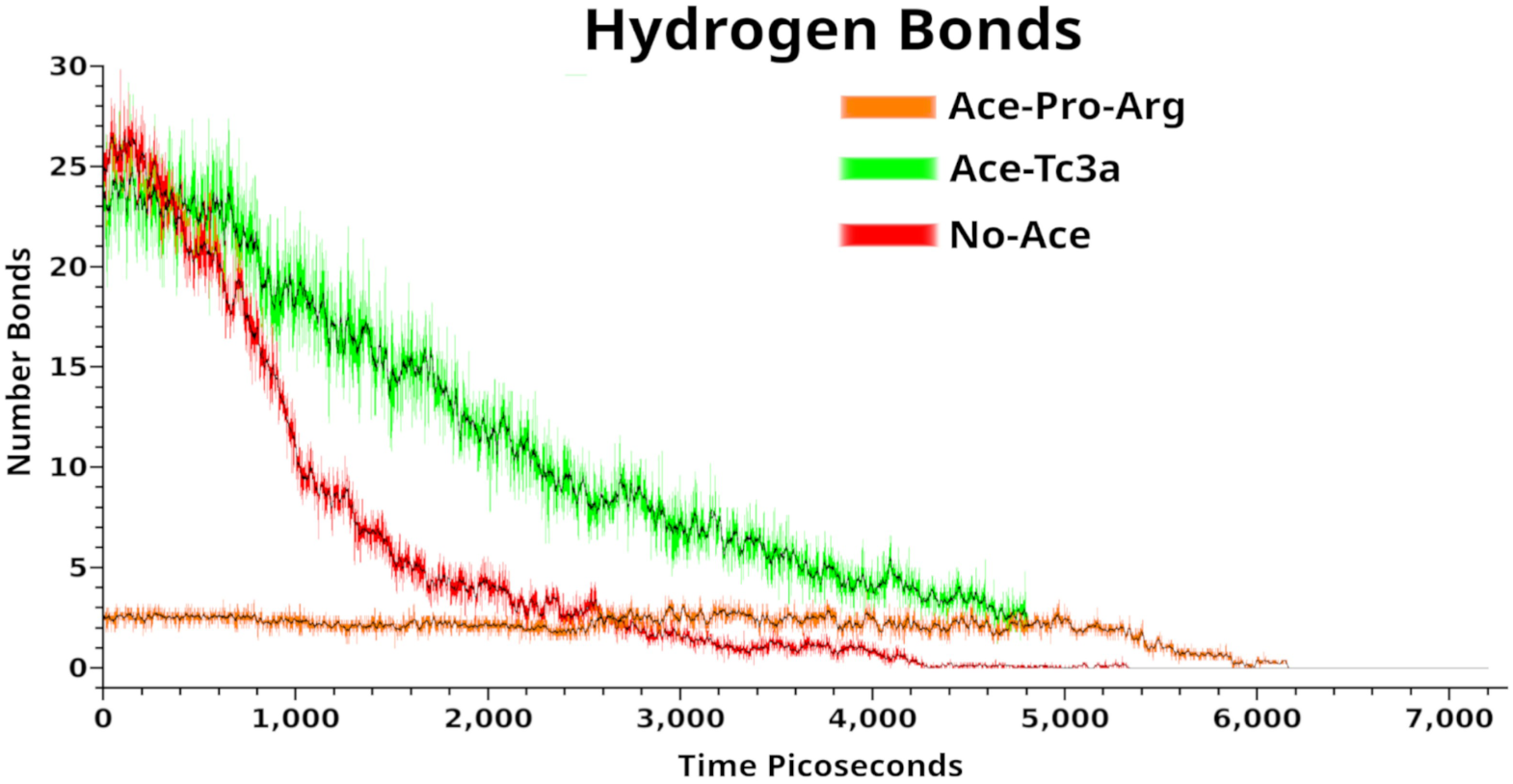
Total NH2-Proline and Acetylated N-terminal Hydrogen Bonding. Hydrogen bonding from pulled simulations as a function of time, green, all hydrogen bonds from Acetylated-Pro-Arg bipartite, red, all hydrogen bonds from NH2-Pro-Arg simulations, orange, only the hydrogen bonds from residues Acetyl-Pro-Arg-Gly N-terminus. Black lines mark the smoothed means of each curve.

We then looked at distances between base pairs atoms, along with distances for the Acetyl group and terminal amino acids. Our general finding was that the DNA stretch from A7 to A12 oscillates between an A form and B form of DNA. When the HTH1 becomes disengaged, the Acetyl-Pro-Arg are the last residues to leave. During these processes, the T13 begins to flip back into a normal conformation, however the force causes A12 and T44 to lose their base pairing. During this process, the NH1, NH2 and NE of Arg3 form transient bonds pulling on A12 and T13 base ends. Mainchain nitrogens of Arg3 and Gly4 then interact with A45 ribose O3’ and O1P, while the H3 of base T13 and N7 of the base ring T45 form a long lasting stable hydrogen bond, Fig. 8. This traps the acetyl group through hydrophobic forces, and with a strong repulsive effect between the oxygen of the group and T45 ribose O4’, as the Acetyl-Pro has no rotational movement between the Pro-Arg mainchain, Fig. 8G,H. This structure is stable for 600-1200 ps in simulations. Interestingly the timing for this processes varied in pulled simulations, and under bulk analysis conditions would be missed with averaged hydrogen bonding, RMSF or spatial analysis techniques. The resulting free energy change for dislodging these interactions is 25-30 k Joules/mol and accounts for the increased affinity, which is determined from the bulk analytical methods employed. None of the other simulations with modified N-terminus formed similar orientations or structures, as the T13 was not unpaired or flipped out in any of these simulations.

**Figure 8.**
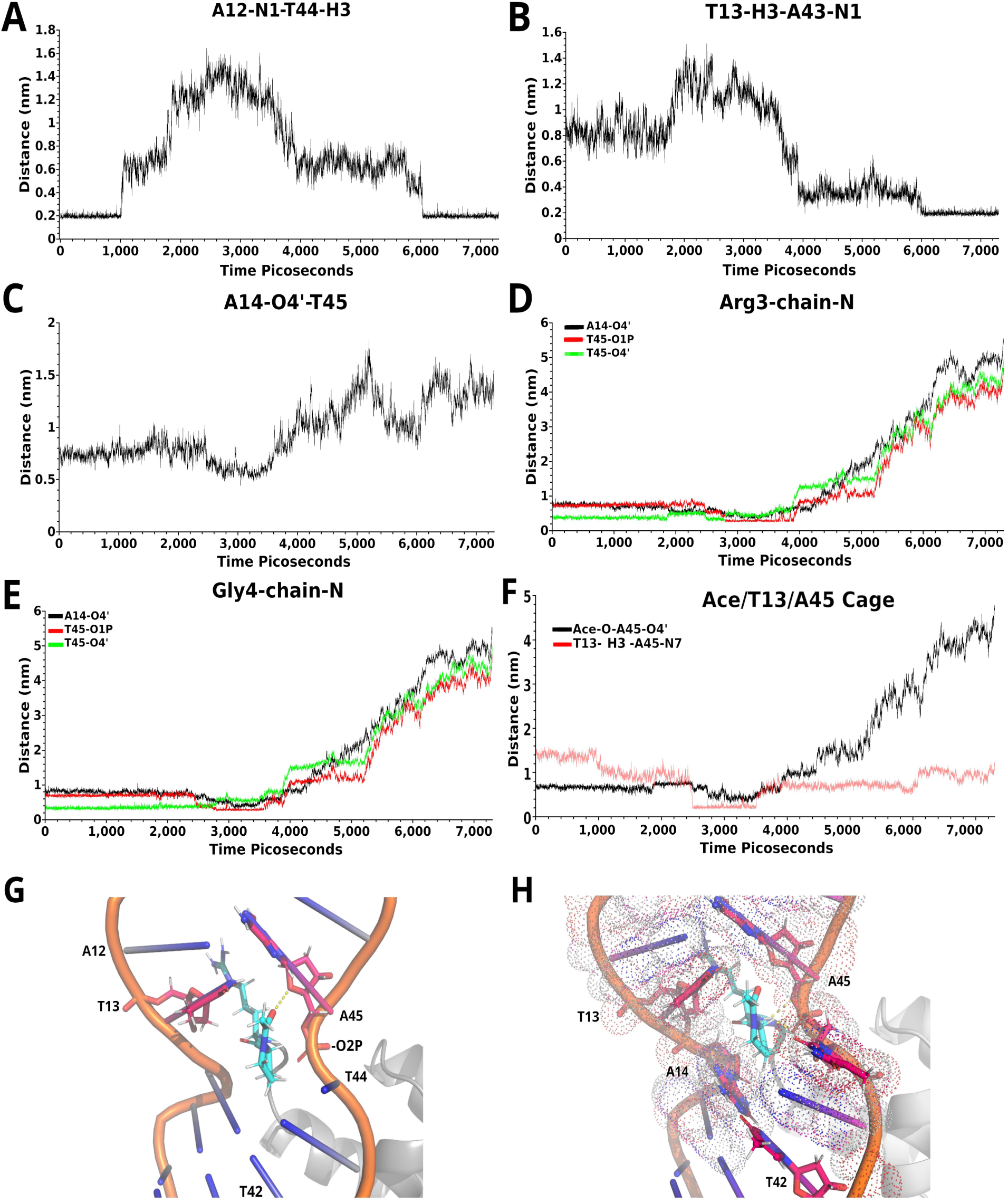
Distance Across Pulled Simulation Between Selected Atoms. A) Base distance between atoms N1 A12 and H3 T44, B) Base distance between atoms H3 T13 and N1 A43, C) Distance between atoms O4’ ribose ring A14 and O1P of T45, D) Composite distances between mainchain NH Arg3 and atoms O4’ ribose ring A14, black, O1P A45, red, and O4’ ribose ring A45, green E) Composite distances between atom NH mainchain Gly4 and atoms O4’ ribose ring A14, black, O1P A45, red, and O4’ ribose ring A45, green F) distances between atom Acetyl O and O4’ ribose ring A45, black, and atom N7 base ring A45 and ring H3 T13, red. G) Structure and visualization of the described intermediate in A-F, snapshot taken at 3000 ps. T42 remains stable while A43 to G46 of the TATAG stretch form a box shape, pinching the phosphates between T13 and A14 together with the phosphate between T44 and A45. The main pinching is between phosphate from T45 and the ribose of A14. Yellow dotted lines mark repulsive interactions under 1.7 angstrom H) Same view as G with dots representing space filling models of atom surfaces. Yellow dotted lines mark hydrogen bonds, T13, A14, T42, T44, A45 represented a stick models, nitrogen blue, hydrogen white, oxygen red, and carbon fuchsia. In both G,H, the Acetyl, Pro2, Arg3 and Gly4 are aqua stick models, nitrogen dark blue, hydrogen white, oxygen red, and carbons aqua. Atom names are those used in the all atom Amber-ILDN force field from simulations.

Utilizing computational proteomics, we attempted to determine if there were any obvious correlations between the processed Acetyl-Pro-Arg N-terminus and DNA binding proteins. We also included dsRNA or RNA/DNA in the analysis, as a structural feature. Using a search algorithm, we extracted all proteins with the respective N-terminus from the Uniprot database, grouping this according to the datasets as presented from source. Using Pfam, Interpro and GO biochemical process identifiers, we then grouped all these into dsDNA and dsRNA binding proteins, S2 Table. Discrepancies exist between Tremble, which includes all sequenced data with or without annotation, and Swiss-protein which has heavily curated and accepted annotated entries. At the organismal level, Plants and Archaea only showed major differences using both dataset types in combination. Bacteria were not included, as they lack acetylation of N-terminal residues, and Archaea only have similar processing of N-terminus in half of sequenced species (66).

To further refine and compare this data, we extracted data sets for H. sapiens only using additionally Met-Pro-His(Ace-PH), Met-Pro-Glu(Ace-PQ), Met-Ser-Arg(Ace-SR), Met-Ala-Leu(Ace-AL) and Met-Arg-Glu(MRQ) for all protein sequences. Then, data was refined to remove alternate spliced isoforms, and only curated proteins which have interactions already known utilized. In some cases only a single isoform with an alternate methionine contains the sequences used, while alternative isoforms have different starting residues. Based on this, we determined all 100% known dsDNA, dsRNA and dsDNA/RNA proteins from the datasets. Statistically, the Acetyl-Pro-Arg was slightly higher than the others, 54.55%, with Ace-PH, Ace-PQ, Ace-SR, Ace-AL at around 30.00% each, and the MRQ which is not processed with methionine removal and acetylation at 7.92%. Enrichment analysis against the human genome with DAVID showed Acetyl-Pro-Arg had 63 proteins out of 126 with a p-value <0.005 that were associated with dsDNA or dsRNA binding, and with MOET p<0.001 again 63, 32 of which were transcription factors and 31 were transcription factor associated. However MRQ showed zero enrichment for any molecular function in MOET, and Ace-AL showed only enrichment for cytokine and interferon receptor associations. Only the acetyl-Ser-Arg in H. sapiens, showed 30.0% of proteins as enriched for direct DNA binding in DAVID, or 35.1% enriched for solely transcription factors in MOET. Tabulated data are presented in S2 Table and Table 2. The python scripts used for basic Uniprot, Tremble, Swiss-protein parsing are available through supplemental materials, and unprocessed raw extracted Uniprot entries listed in S1.File.

**Table 2.** Comparison of different H.sapiens N-terminal extracted proteins from Uniprot datasets.

| N-terminus | Number Fully Annotated | Number dsDNA/ RNA binding | DAVID Percent Enrichment dsDNA/RNA interacting | MOET Percent Enrichment |
| --- | --- | --- | --- | --- |
| Ace-PR | 112 | 61 | 54.55% | 50% DNA transcription/ 8% DNA editing/ 23% RNA binding |
| Ace-PH | 105 | 36 | 34.29% | 50% Oxoreductase, 35% single nucleotide binding, 20% lipid oxidizing, 5% transcription factor |
| Ace-PQ | 108 | 32 | 29.63% | 23% α-actin associated |
| Ace-SR | 150 | 45 | 30.00% | 35.10% transcription factor |
| Ace-AL | 344 | 73 | 21.22% | 48.5% Cytokine receptor associated, 20% interferon receptor associated |
| MRQ | 101 | 8 | 7.92% | 0% enrichment |

The N-terminus used is listed to the left first column. Only MRQ is not processed by human processing machinery into acetylated, methionine cleaved N-terminus. Column 2 is the number of proteins completely annotated with the respective N-terminus. Column 3 is the number of proteins that have direct dsDNA, dsRNA or dsDNA/RNA interactions listed in their functional identifier entries directly. Both column 2 and 3 manually checked to include identifier. Column 4 is the percentage shown to be enriched for double stranded nucleotide interacting processed through DAVID, with p<0.005 acceptance. Column 5 is the entire Uniprot H.sapiens Tremble and Swiss Protein datasets with all alternate spliced isoforms, and redundant entries removed, then parsed through MOET to identify functional enrichment, shown as percent of total included proteins, (122 ace-Pro-His min to 1117 ace-Ala-Leu max), with p-values <0.001. DAVID contains more broad nucleic acid definitions, while MOET has more precisely binned identifiers for functionality.

## Discussion

Our data clearly shows the acetylated processed N-terminus plays an important role in the Tc3a transposition process. Here the effect is approximately 77 k Joules higher affinity compared with the NH2-Pro-Arg alone showing the difference cannot be attributed to hydrogen bonding, with only a single additional oxygen. The processes involved rather result from drastic differences in DNA structure induced by the different N-terminal structure highlighting the sharp effects of small molecular changes. Acetyl groups only have 6 atoms, with only 1 charged oxygen and in the Tc3a N-terminus attach directly to the proline terminal nitrogen. This resulting change causes a flipping of T13 in the TATA region of the Tc3a ITR, which is not mirrored in other N-terminus tested here, including the addition of an unprocessed methionine. Higher affinity is the result of this induced DNA conformation, and the effects it has on the DNA during the binding and dissociation processes of the Tc3a bipartite, as presented in Fig. 2 and Fig. 8. A novel stable DNA-protein intermediate is responsible for 25-30 k Joule differences observed from all tested N-terminus in the study, which occurs for an extended period in molecular time frames between 600-1200 ps. This is enough time for the HTH1 domain of Tc3a to dissociate and re-associate in natural conditions, without an applied force. Additional higher affinity is achieved by the changes in the minor groove, particularly with the loop region binding, also induced by the acetyl group.

Base flipping has been observed in other transposon families, outside of mariners, and has been proposed to be a common mechanism in many DNA binding protein’s function (67–69). Base flipping alone has been characterized for double stranded DNA in the past, and shown to generate a force on DNA equal to a right handed vector, moving in the 3’ to 5’ direction of the strand involved from published literature (70–73). While our experimental set up can not determine an accurate empirical force, which would require 2-3x longer DNA in simulations, and extended simulations of the DNA alone, we are able to determine the direction of propagation and effects from the bipartite on the DNA. Our spatial analysis shows the HTH2 acts to dampen any effects from the flipped base, and changes motions mainly into an x-y direction. This then induces a widened stretched out base pairing that is held in a stable conformation, with little movement along the length of DNA. Contrastingly, the motion of propagation is then transferred to the HTH2 direction, which coincides with the direction of the opposite ITR of the transposon.

In Tc3a or similar related transposase, the process most likely plays a role in the initial ITR cleavage process, which has several competing proposed mechanisms (19,20,74,75). Currently these include supercoiling to bring single transposase together, dimerization at one or both ends, and a multimerization of 4 single mariners which in some way generate torsion to bring the ITR ends of the transposon together. Torsion provided by the single base flipping at either ITR would produce enough force for supercoiling of dsDNA, where the 1200 base pairs between would wind around itself bringing the two ends together. In published work with extended DNA, base flipping has been shown to supercoil stretches between two flipped bases, and conversely to result from forced supercoiling (72–76). Of the debated mechanisms with supercoiling, one hypothesis includes a single nick at either end and bending of the ITR cause a similar process, however base flipping has never been observed in Tc3a or Mariners. Much of these differences in theoretical processes may be an artifact of the expression systems used, with all X-ray and NMR structures published utilizing E. coli which lacks the acetylation processing. It has also been mentioned in literature that the HTH1, or bipartite domains of mariners have lower than expected affinities in DNA binding wet laboratory experiments, which utilize the same expression systems and are routinely used in most wet laboratory experiments to develop all process models.

Our deduced process however may be Tc3a specific, with only 4% of known mariner transposons having an N-terminal acetylated Pro-Arg, although this number increases slightly if Mos1, piggybac, or related mariner splice variants are included to 10%. Also a number of transgenic mariner evolved fusion proteins such as SETMAR in humans and Papio anubis retain the bipartite domain at the C-terminus without the acetylated end or catalytic domain. However with SETMAR and others there are maintained isoforms where the entire expressed fusion protein still retains an acetylated PR N-terminus (76–78). In the PAX proteins, proposed to be evolutionary related proteins to mariners from horizontal transfer, only one from a dozen maintains a acetyl-PR N-terminus, although others have low expressed isoforms with the acetyl-PR or PH. However almost the entire groups from transcription factor families FOXB, TSZH and GSX1 have acetyl-PR N-terminus. Interestingly Phytophthora and Planariidae have a larger portion of identified mariner transposase with this Acetyl-PR N-terminus, grouped into Tc3 like or Tc1 like. Still, without further sub-grouping transposase, mariner transposase must have alternate mechanisms for the same process if base flipping is intrinsic to the family as a whole. It has been shown in Tn5 and Tn10 transposase, which utilize different mechanisms including hairpin formations, base flipping 3-4 bases from the ITR cleavage site is mediated by forcing a methionine between the bases, aided by the bound transposase. Similar processes to the one described in our work may be possible with alternative sequences, which would include 8000 novel variants if all 3 primary amino acids are included. Ultimately, our work indicates that N-terminal processing needs to be taken into account in the study of DNA binding proteins in general.

More theoretical, we attempted to find a significant correlation with the Acetyl-PR N-terminus and dsDNA, dsRNA or dsDNA/RNA binding proteins. Our data indicates a slightly higher correlation for the N-terminal residues analyzed, however the increase of only 20-25% more from others indicates this may be in a more specific protein domain or family aspect. In simple enrichment analysis, the Acetyl-PR did show high enrichment over other tested N-terminal sets, however when looking at the protein sets individually all contained DNA binding proteins. Additionally enrichment analysis routinely left out proteins listed as direct dsDNA or dsRNA binding in their descriptions. In a number of mammalian mariner evolved fusion proteins, the HTH1 and HTH2 are on the C-terminus, while the Acetyl-PR still make up the N-terminal domain, and are in direct contact with the DNA, suggesting this still may have some role. We were however hindered from direct structural analysis as only 8 structures from the 476 human proteins, including splice variants identified, S2 Table and Tabel 2, had structures complete enough to analyzed. From these 3 showed the methylated end formed a β-sheet that was twisted at the ends in the X-ray structures lacking processed N-terminus, 3 were ribosomal structural proteins embedding the N-terminus in dsRNA and lacking the acetyl group, 1 was a ribonuclear protein with dsDNA/RNA, and 1 a transcription factor superposable with Tc3a again lacking the acetyl group. This sample size is to small to make direct conclusions or provide statistic on function, making our proteomics analysis only suggestive for common functional categorizations.

## Conclusion

Our work here shows acetylation in N-trminal protein processing plays a role in part of the Tc3a transposase transposition process, conferring higher affinity and altering the DNA structure. The overall processes of Mariner transposition, presented in the introduction, are outlined in Fig. 9. For Tc3a specifically, the flipped base at either end of the transposon would provide force to cause the DNA between the sites to supercoil. This supports a model outlined by other groups, where single transposase bind at either end and are brought together to form a dimer from DNA torsion alone, outlined in Fig. 9, 1 and 4. A model similar to this has been proposed as a general mechanism used to bring together distant transcription factors or DNA binding proteins in larger complexes (16,21,68,76).

**Figure 9.**
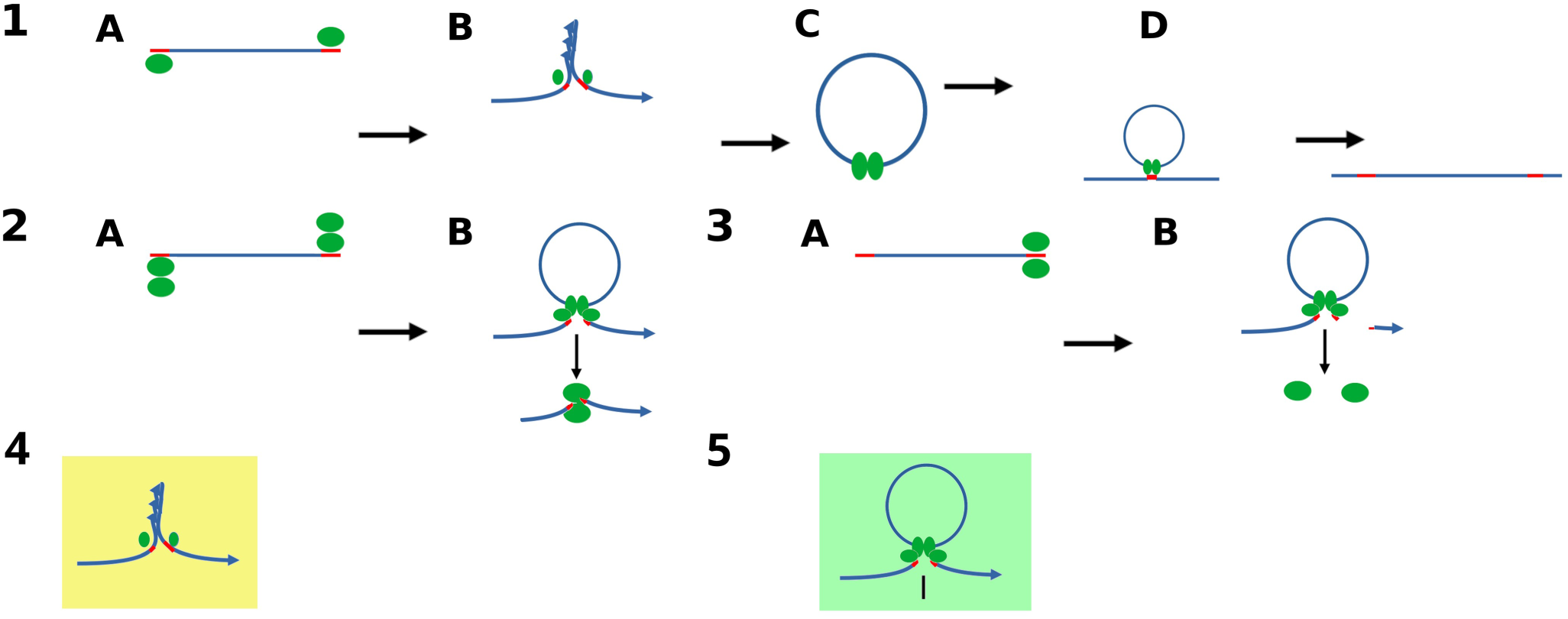
Proposed Tc3a/Mariner excision process. 1) A) A single transposase binds the ITR of the transposon at either end. B) Distortion on the DNA at either end causes the region between ITR to super-coil, bringing the two transposase together, allowing for the cleavage of both strands at the ITR end. C) The dimerized complex leaves the excision site, which is repaired by DNA ligase leaving a TATA duplication, D) remaining dimerized the complex then scans DNA until a TA site is recognized, inserts the transposon complete with the ITR at either end, and the two proteins dissociate. C and D remain the same for all models, except the coiling or looped DNA are both proposed in different models to make up the complex until insertion. 2) A) Dimerized transposase binds either end of the transposon at the ITR, B) These cause torsion that brings the 4 transposase together, which then are able to cleave both strands of DNA, one of each original dimer leaves with the excision site, while the other leaves carrying the transposon. 3) A) Two transposase bind one ITR of the transposon B) these dimerize with unbound transposase forming a 4 protein complex excising one end of the transposon, and coil the DNA until the second ITR is found, and the entire transposon is cleaved at either ITR. This leaves the two excised ends distant from each other. The excision site at a lower frequency has 1-4 base pairs making a TAX_N_TA marker, which has been statistically analized for Tc3a in C.elegans. Variations exist on these 3 main proposed mechanisms. Green ovals, transposase, red lines, ITR, blue lines, dsDNA, in 3B, half red line a TA overhang. 4,5) Representatives of the two main remaining questions in Mariner transposition processes; Do the transposase form multimeric complexes to cleave the DNA, or do they work monomeric, and how are the two ends of the transposon brought together, passively or through DNA induced torsion?

Still, this does not address the order of DNA strand cleavage, which still contains models of single strand or double strand nicking also playing a role in bringing two transposase together. However, it does indicate that multimerization at either ITR end is not necessary for the initial inverted terminal repeat torsions necessary for the overall process. Thus more complex 4 transposase multimeric systems are most likely not necessary in mariner ITR dimerization processes. We propose a more simplistic system based on the DNA binding from processed terminal residues and induced torsions in the ITR as a major factor driving dimerization, and the entire process of excision.

Supplementary Data are available in supplementary materials for download, models used are deposited on the ModelArchive at https://www.modelarchive.org/10.5452/ma-5m69p

Python scripts used for Uniprot parsing are available at https://github.com/StephanWatkins/Uniprot-Python-parsing

## AUTHOR CONTRIBUTIONS

Helder I. Nakaya:Conceptualization, Methodology,Validation,Writing-review&editing, Resources, Funding acquisition

Stephan L. Watkins:Conceptualization, Methodology,Validation, Formal analysis, Writing

## ACKNOWLEDGMENTS

Use of grid computers to conduct initial and subsequent simulations was made possible through: Laboratório Nacional de Computação Científica - Av. Getúlio Vargas, 333, Quitandinha, Petrópolis, RJ, Brasil, CEP 25651-075 and CENAPAD-SP Rua Saturnino de Brito, 45, Cidade Universitária, Campinas, SP, Brasil, CEP 13083-889.

## FUNDING

Change to Financial disclosure: “This work was supported by visiting researcher grant at the Institute Israelite Teaching and Research Albert Einstein (IIEPAE): Brazilian Beneficent Society Israelite, Hospital Albert Einstein (SBIBAE, grant number CNPJ 11.445.958/0001-02). Helder I. Nakaya received funding for this work through FAPESP (Fundação de Amparo à Pesquisa do Estado de São Paulo, grant number: 2018/14933-2) and CNPq (Conselho Nacional de Desenvolvimento Científico e Tecnológico, grant number 313662/2017-7). The funders had no role in study design, data collection and analysis, or preparation of the manuscript. Decision to publish is mandatory under the granting guidelines.

## CONFLICT OF INTEREST

Helder I. Nakaya is the Chief Scientific Officer (CSO) of Synamics Therapeutics, a startup that utilizes artificial intelligence for drug discovery. This does not alter our adherence to PLOS ONE policies on sharing data and materials.

---

**Supplemental Figure 1.**
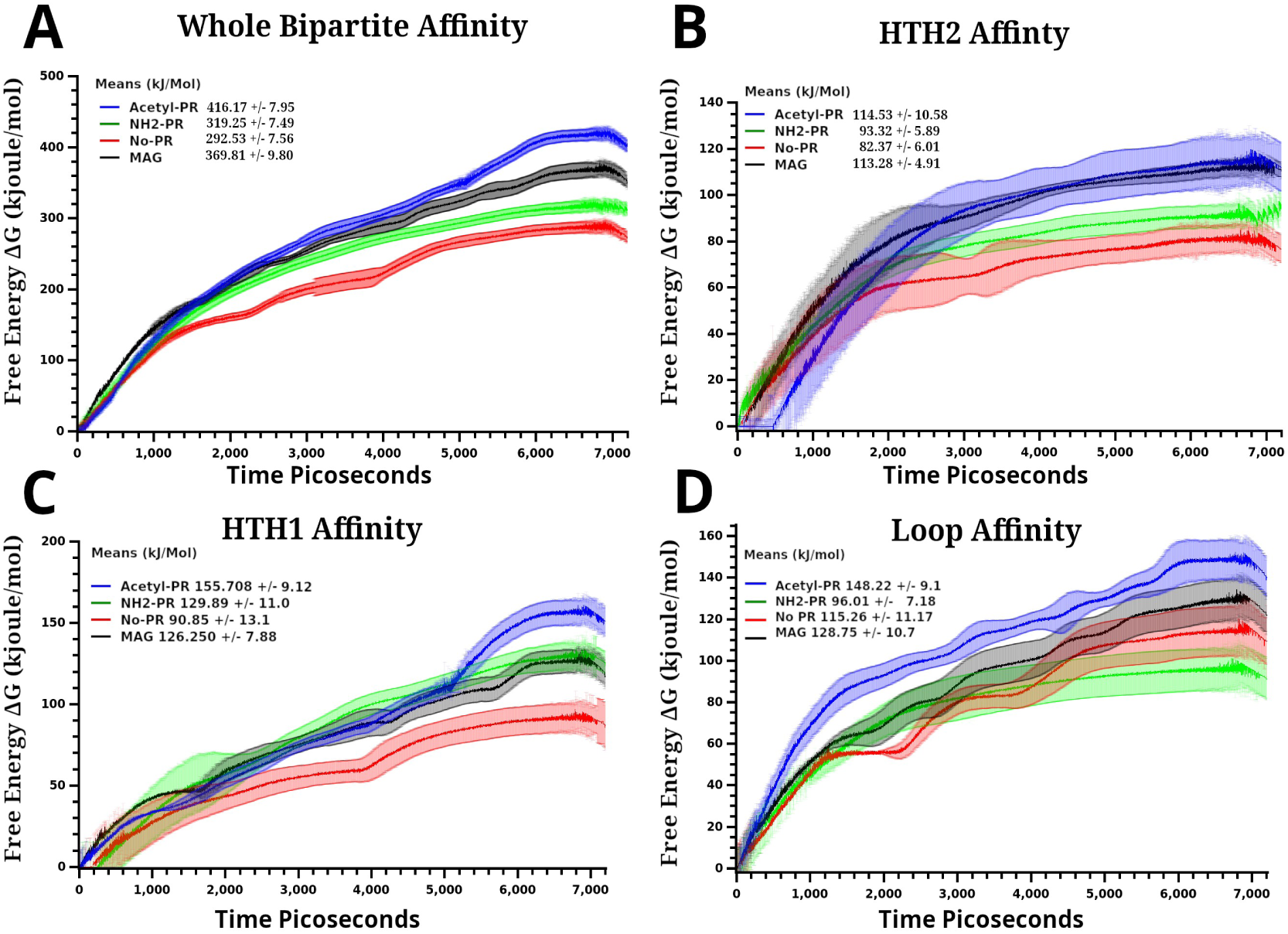
Calculated Total and Bipartite Domain Affinities with Standard Deviation. Figure are the same as Figure 3, with all standard deviation included. A) Binding affinity in kJoules/mol, and Gibbs free energy (ΔG) change of the entire bipartite domain averaged from all pulled simulations versus time, 7.2 ns, B) calculated mean affinity, and ΔG for HTH2, C) calculated mean affinity and ΔG for HTH1 , D) Binding affinity means for Loop region, residues Val47-Lys59, and ΔG over time. Standard deviation are given as +/-either side of mean for A-D mean affinities. In all, Acetylated PR, blue, MAG, black, NH2-PR, green, and truncated to NH2-G end, red. Standard deviation are given as +/-either side of mean for A-D across runs, and are shaded the same color as the respective legend line.

**Supplemental Figure 2.**
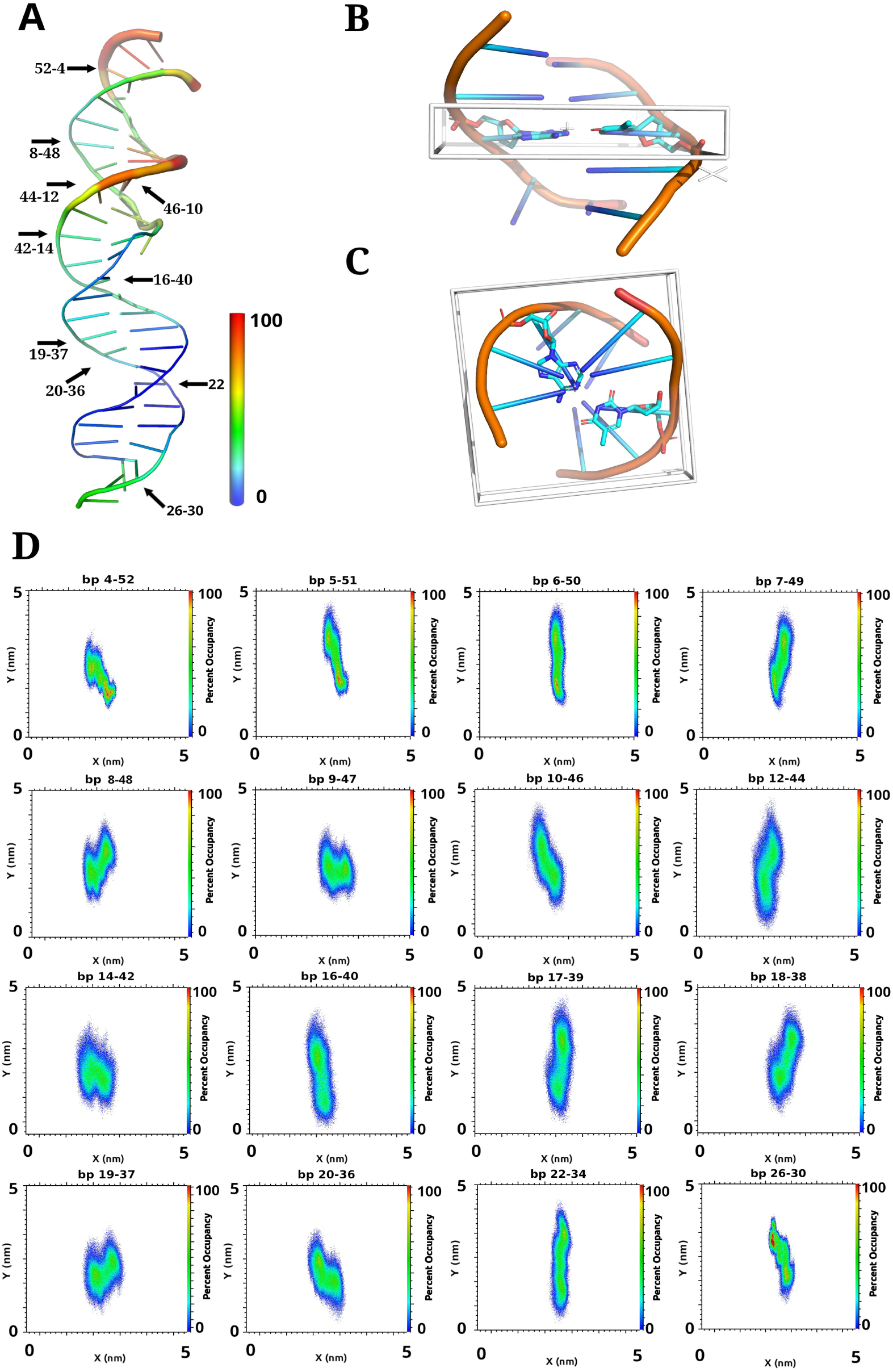
Occupied Density Maps of Base Pairs During Extended Simulations. **A)** Jelly role with RMSF standardized B-factors for bound ITR DNA from 50ns of extended simulation time. Arrows indicate respective base pair densities in D), **B)** Side view and **C)** top of a representative region used for generating the density plots in **D)**. The 6 point white star in B) represents the center axis, which extended through the short axis of the box as Z, box white, marking the density region. The box extends to the periodic boundaries in the X and Y direction, and the base pair densities are centered in the region as close as possible, with a 5nm x 5nm box size . Plots are projected onto an X-Y plane, with total density represented as a single X and Y vector in nm^3^. The total plot area represent total space occupied for the simulation, D) Base pairs are listed above density plots, density is scaled as 0.1%-100% occupancy according to color scales on the right of each representation, converted to the maximum density peaks in a 0.001 x 0.001 nm 2D area set to 18 nm^3^ , white, 0% occupancy. Base pair 4-52 represents a stable normal X-Y fluctuating B-DNA pair for reference, despite high RMSF in A) the base pairs form a classical compact interaction. All represent the total occupancy from all frames of 50ns simulation time, centered between pair phosphates in the x-y direction, and +/-3.5 Å in the Z direction.

**Supplemental Table 1.** Comparison of X-ray and Molecular Dynamics Structures

| X-ray |  |  | MD |
| --- | --- | --- | --- |
|  |  | Missing in MD | Missing in X-ray |
| Side Chain |  |  | Side chain |
| Protein | DNA |  | Protein |
| Arg3-NE | T11-O2 |  | Cys38 |
| Ser24-OG | G15-OP1 |  | Arg40 |
| His26-NE | G6-N7 |  | Ser52 |
| Arg30-NH2 | G5-OP2 |  | Lys53 |
| Arg34-NH1 | G46-OP1 |  | Arg58 |
| Ser35-OG | G46-OP2 |  | Lys59 |
| Arg36-NH1 | G7-N7 |  | Thr79 |
| Arg36-NH2 | G7-O6 |  | Arg81 |
| His37-ND1 | G46-N7 |  | Lys101 |
| Tyr49-OH | A45-OP1 | xx |  |
| Arg54-NE | T42-O2 |  |  |
| Arg54-NH2 | A43-N3 |  |  |
| Arg57-NH1 | T39-O2 |  |  |
| Ser92-OG | T20-OP2 |  |  |
| Lys93-NZ | G32-N7 |  |  |
| Arg94-NE | T42-O2 |  |  |
| Arg94-NH2 | G34-N7 |  |  |
| Thr95-OG1 | T20-OP2 |  |  |
| Arg102-NH2 | T19-OP2 |  |  |
| Backbone |  |  | Backbone |
| Pro2-N | T13-O2 | xx | Gly50 |
| Arg3-N | A54-N3 |  | Thr51 |
| Gly4-N | A45-O4' |  | Ser52 |
| Ala6-N | G15-OP1 | xx | Ala55 |
| Leu25-N | G6-OP1 |  | Pro56 |
| His26-N | G6-OP2 |  | Arg57 |
| Ser35-N | G46-OP2 |  | Lys59 |
| Arg58-N | C41-OP1 |  | ALA91 |
| Ala60-N | T19-OP1 |  |  |
| Ala80-N | G32-OP1 |  |  |
| Ser92-N | T20-OP2 |  |  |

Columns 1 and 2 list hydrogen bond pairs found in the X-ray structure. These are separated first into amino acid or DNA as column 1 and 2, then into side chain or backbone interactions, top to bottom. Column 3 the xx denote hydrogen bonds not found in the molecular simulations. Column 4 is a list of all hydrogen bonds not found, from the respective amino acids, in the X-ray structure. The lifetimes and atoms involved are listed in table 1.

**Supplemental Table 2.** Tabulated Uniprot Parsed N-terminal Acetyl-Pro-Arg Proteins

| Uniprot Dataset | Number Entries | DNA Associated | RNA-Associated |
| --- | --- | --- | --- |
| H.sapiens |  |  |  |
| Tremble | 358 | 37.70% | 40.20% |
| Swiss-prot | 129 | 69.80% | 67.40% |
| Rodent |  |  |  |
| Tremble | 4546 | 50.70% | 47.60% |
| Swiss-prot | 229 | 60.30% | 60.70% |
| Mammals |  |  |  |
| Tremble | 25416 | 41.80% | 42.00% |
| Swiss-prot | 74 | 40.50% | 52.70% |
| Vertebrates |  |  |  |
| Tremble | 11858 | 39.30% | 40.00% |
| Swiss-prot | 198 | 60.10% | 59.10% |
| Invertebrates |  |  |  |
| Tremble | 52472 | 27.70% | 28.60% |
| Swiss-prot | 89 | 61.80% | 77.50% |
| Plants |  |  |  |
| Tremble | 48196 | 19.90% | 23.80% |
| Swiss-prot | 156 | 22.40% | 28.20% |
| Fungi |  |  |  |
| Tremble | 109083 | 27.80% | 32.70% |
| Swiss-prot | 154 | 63.60% | 74.00% |
| Archaea |  |  |  |
| Tremble | 23518 | 20.80% | 29.30% |
| Swiss-prot | 104 | 43.30% | 76.90% |
| Virus |  |  |  |
| Tremble | 9827 | 36.40% | 36.90% |
| Swiss-prot | 325 | 63.70% | 62.70% |

Number of proteins, grouped according to Uniprot datasets found with N-terminal Acetyl-Pro-Arg. Percentages are all proteins found with GO function, Pfam, and Interpro composited for DNA or RNA interacting. Overlap exists between DNA and RNA interacting as a number of proteins are also dsDNA/RNA interacting or ribonuclear proteins that interact with dsDNA. A number of Tremble and a small portion of Swiss protein (~10-15% and 1-2% respectively) have unknown functions. Column 1, organism division, and Tremble or Swiss protein dataset, column 2, number of entries found, column 3, percent with dsDNA interaction identifiers, column 4, percent with dsRNA or ribonuclear protein identifiers.

## Supplemental Movies

**Sup.Video1-** NH2-Pro-Arg

**Sup.Video2-** Acetyl-Pro-Arg, acetyl black

**Sup.Video3-** No Pro-Arg3

**Sup.Video4-** MAG

All videos are oriented as in Figure 6B, and aligned before generating the video images.

## Supplemental File

List of Uniprot IDs extracted from the Human Uniprot Trembel and Swiss-Protein datasets.

## Notes

### Summary of Updates

Initial review process for publication. All changes were to address comments, concerns or issues reviewers or editorial staff made to move towards publication.

## References

1. Mc CB. Maize genetics. Year B Carnegie Inst Wash. 1946;45:176–86.

2. Mc CB. The origin and behavior of mutable loci in maize. Proc Natl Acad Sci U S A. 1950 Jun;36(6):344–55.

3. Feschotte C. Transposable elements and the evolution of regulatory networks. Nat Rev Genet. 2008 May;9(5):397–405.

4. Plasterk RHA, van Luenen H. Transposons. In: Riddle DL, Blumenthal T, Meyer BJ, Priess JR, editors. C elegans II. Cold Spring Harbor (NY): Cold Spring Harbor Laboratory Press Copyright © 1997, Cold Spring Harbor Laboratory Press.; 1997.

5. Feschotte C, Pritham EJ. DNA transposons and the evolution of eukaryotic genomes. Annu Rev Genet. 2007;41:331–68.

6. Wang S, Diaby M, Puzakov M, Ullah N, Wang Y, Danley P, et al. Divergent evolution profiles of DD37D and DD39D families of Tc1/mariner transposons in eukaryotes. Mol Phylogenet Evol. 2021 Aug;161:107143.

7. Gao B, Wang Y, Diaby M, Zong W, Shen D, Wang S, et al. Evolution of pogo, a separate superfamily of IS630-Tc1-mariner transposons, revealing recurrent domestication events in vertebrates. Mob DNA. 2020;11:25.

8. Wang W, Swevers L, Iatrou K. Mariner (Mos1) transposase and genomic integration of foreign gene sequences in Bombyx mori cells. Insect Mol Biol. 2000 Apr;9(2):145–55.

9. Yokoo M, Fujita R, Nakajima Y, Yoshimizu M, Kasai H, Asano S, et al. Mos1 transposon-based transformation of fish cell lines using baculoviral vectors. Biochem Biophys Res Commun. 2013 Sep 13;439(1):18–22.

10. Woodard LE, Wilson MH. piggyBac-ing models and new therapeutic strategies. Trends Biotechnol. 2015 Sep;33(9):525–33.

11. Kim SW, Lee JH, Han JS, Shin SP, Park TS. piggyBac Transposition and the Expression of Human Cystatin C in Transgenic Chickens. Animals (Basel). 2021 May 26;11(6).

12. Plasterk RH. The Tc1/mariner transposon family. Curr Top Microbiol Immunol. 1996;204:125–43.

13. Wells JN, Feschotte C. A Field Guide to Eukaryotic Transposable Elements. Annu Rev Genet. 2020 Nov 23;54:539–61.

14. Pace JK 2nd, Feschotte C. The evolutionary history of human DNA transposons: evidence for intense activity in the primate lineage. Genome Res. 2007 Apr;17(4):422–32.

15. Craig NL. Target site selection in transposition. Annu Rev Biochem. 1997;66:437– 74.

16. Morris ER, Grey H, McKenzie G, Jones AC, Richardson JM. A bend, flip and trap mechanism for transposon integration. Sherratt D, editor. eLife. 2016 May;5:e15537.

17. Galbraith JD, Ivancevic AM, Qu Z, Adelson DL. Detecting Horizontal Transfer of Transposons. Methods Mol Biol. 2023;2607:45–62.

18. Palazzo A, Escuder E, D’Addabbo P, Lovero D, Marsano RM. A genomic survey of Tc1-mariner transposons in nematodes suggests extensive horizontal transposon transfer events. Mol Phylogenet Evol. 2021 May;158:107090.

19. Ochmann MT, Ivics Z. Jumping Ahead with Sleeping Beauty: Mechanistic Insights into Cut-and-Paste Transposition. Viruses. 2021;13(1).

20. Claeys Bouuaert C, Chalmers R. A single active site in the mariner transposase cleaves DNA strands of opposite polarity. Nucleic Acids Research. 2017 Nov 16;45(20):11467–78.

21. Tellier M, Bouuaert CC, Chalmers R. Mariner and the ITm Superfamily of Transposons. Microbiol Spectr. 2015 Apr;3(2):MDNA3-0033-2014.

22. Jaillet J, Genty M, Cambefort J, Rouault JD, Augé-Gouillou C. Regulation of mariner transposition: the peculiar case of Mos1. PLoS One. 2012;7(8):e43365.

23. Richardson JM, Colloms SD, Finnegan DJ, Walkinshaw MD. Molecular architecture of the Mos1 paired-end complex: the structural basis of DNA transposition in a eukaryote. Cell. 2009 Sep 18;138(6):1096–108.

24. Hickman AB, Dyda F. Mechanisms of DNA Transposition. Microbiol Spectr. 2015 Apr;3(2):MDNA3-0034–2014.

25. Zhou L, Mitra R, Atkinson PW, Hickman AB, Dyda F, Craig NL. Transposition of hAT elements links transposable elements and V(D)J recombination. Nature. 2004 Dec 23;432(7020):995–1001.

26. Heyer WD, Ehmsen KT, Solinger JA. Holliday junctions in the eukaryotic nucleus: resolution in sight? Trends Biochem Sci. 2003 Oct;28(10):548–57.

27. Harshey RM. Transposable Phage Mu. Microbiol Spectr. 2014 Oct;2(5).

28. Dupeyron M, Baril T, Bass C, Hayward A. Phylogenetic analysis of the Tc1/mariner superfamily reveals the unexplored diversity of pogo-like elements. Mobile DNA. 2020 Jun 29;11(1):21.

29. Kojima KK. Structural and sequence diversity of eukaryotic transposable elements. Genes Genet Syst. 2020 Jan 30;94(6):233–52.

30. Watkins S, van Pouderoyen G, Sixma TK. Structural analysis of the bipartite DNA-binding domain of Tc3 transposase bound to transposon DNA. Nucleic Acids Res. 2004;32(14):4306–12.

31. Wingfield PT. N-Terminal Methionine Processing. Curr Protoc Protein Sci. 2017 Apr 3;88:6.14.1-6.14.3.

32. Van Der Spoel D, Lindahl E, Hess B, Groenhof G, Mark AE, Berendsen HJ. GROMACS: fast, flexible, and free. J Comput Chem. 2005 Dec;26(16):1701–18.

33. Darden T, Perera L, Li L, Pedersen L. New tricks for modelers from the crystallography toolkit: the particle mesh Ewald algorithm and its use in nucleic acid simulations. Structure. 1999 Mar 15;7(3):R55–60.

34. Humphrey W, Dalke A, Schulten K. VMD: visual molecular dynamics. J Mol Graph. 1996 Feb;14(1):33–8, 27–8.

35. Jurrus E, Engel D, Star K, Monson K, Brandi J, Felberg LE, et al. Improvements to the APBS biomolecular solvation software suite. Protein Sci. 2018 Jan;27(1):112– 28.

36. Baker NA, Sept D, Joseph S, Holst MJ, McCammon JA. Electrostatics of nanosystems: application to microtubules and the ribosome. Proc Natl Acad Sci U S A. 2001 Aug 28;98(18):10037–41.

37. Dolinsky TJ, Nielsen JE, McCammon JA, Baker NA. PDB2PQR: an automated pipeline for the setup of Poisson-Boltzmann electrostatics calculations. Nucleic Acids Res. 2004 Jul 1;32(Web Server issue):W665–667.

38. Lill MA, Danielson ML. Computer-aided drug design platform using PyMOL. J Comput Aided Mol Des. 2011 Jan;25(1):13–9.

39. Makarewicz T, Kaźmierkiewicz R. Molecular dynamics simulation by GROMACS using GUI plugin for PyMOL. J Chem Inf Model. 2013 May 24;53(5):1229–34.

40. Hub JS, de Groot BL, van der Spoel D. g_wham:A Free Weighted Histogram Analysis Implementation Including Robust Error and Autocorrelation Estimates. Journal of chemical theory and computation. 2010;6(12):3713–20.

41. Lemkul JA. Introductory Tutorials for Simulating Protein Dynamics with GROMACS. J Phys Chem B. 2024 Oct 3;128(39):9418–35.

42. Elber R, Ruymgaart AP, Hess B. SHAKE parallelization. Eur Phys J Spec Top. 2011 Nov 1;200(1):211–23.

43. Luzar A, Chandler D. Effect of environment on hydrogen bond dynamics in liquid water. Phys Rev Lett. 1996 Feb 5;76(6):928–31.

44. Lindorff-Larsen K, Piana S, Palmo K, Maragakis P, Klepeis JL, Dror RO, et al. Improved side-chain torsion potentials for the Amber ff99SB protein force field. Proteins. 2010 Jun;78(8):1950–8.

45. Briones R, Blau C, Kutzner C, de Groot BL, Aponte-Santamaría C. GROmaρs: A GROMACS-Based Toolset to Analyze Density Maps Derived from Molecular Dynamics Simulations. Biophys J. 2019 Jan 8;116(1):4–11.

46. Kuzmanic A, Zagrovic B. Determination of ensemble-average pairwise root mean-square deviation from experimental B-factors. Biophys J. 2010 Mar 3;98(5):861–71.

47. Ahamad S, Hema K, Ahmad S, Kumar V, Gupta D. Insights into the structure and dynamics of SARS-CoV-2 spike glycoprotein double mutant L452R-E484Q. 3 Biotech. 2022 Apr;12(4):87.

48. Dixit SB, Mezei M, Beveridge DL. Studies of base pair sequence effects on DNA solvation based on all-atom molecular dynamics simulations. J Biosci. 2012 Jul;37(3):399–421.

49. Wellenzohn B, Flader W, Winger RH, Hallbrucker A, Mayer E, Liedl KR. Influence of netropsin’s charges on the minor groove width of d(CGCGAATTCGCG)2. Biopolymers. 2001 2002;61(4):276–86.

50. Fratini AV, Kopka ML, Drew HR, Dickerson RE. Reversible bending and helix geometry in a B-DNA dodecamer: CGCGAATTBrCGCG. J Biol Chem. 1982 Dec 25;257(24):14686–707.

51. Laughton C, Luisi B. The mechanics of minor groove width variation in DNA, and its implications for the accommodation of ligands. J Mol Biol. 1999 May 21;288(5):953–63.

52. Crooks GE, Hon G, Chandonia JM, Brenner SE. WebLogo: a sequence logo generator. Genome Res. 2004 Jun;14(6):1188–90.

53. The UniProt Consortium. UniProt: the Universal Protein Knowledgebase in 2025. Nucleic Acids Research. 2024 Nov 18;53(D1):D609–17.

54. Paysan-Lafosse T, Andreeva A, Blum M, Chuguransky SR, Grego T, Pinto BL, et al. The Pfam protein families database: embracing AI/ML. Nucleic Acids Res. 2024 Nov;53(D1):D523–34.

55. Blum M, Andreeva A, Florentino LC, Chuguransky SR, Grego T, Hobbs E, et al. InterPro: the protein sequence classification resource in 2025. Nucleic Acids Research. 2024 Nov 20;53(D1):D44–456.

56. Thomas PD, Ebert D, Muruganujan A, Mushayahama T, Albou LP, Mi H. PANTHER: Making genome-scale phylogenetics accessible to all. Protein Science. 2022 Jan 1;31(1):8–22.

57. Vedi M, Nalabolu HS, Lin CW, Hoffman MJ, Smith JR, Brodie K, et al. MOET: a web-based gene set enrichment tool at the Rat Genome Database for multiontology and multispecies analyses. Genetics. 2022 Apr 4;220(4).

58. Sherman BT, Hao M, Qiu J, Jiao X, Baseler MW, Lane HC, et al. DAVID: a web server for functional enrichment analysis and functional annotation of gene lists (2021 update). Nucleic Acids Res. 2022 Jul 5;50(W1):W216–w221.

59. van Pouderoyen G, Ketting RF, Perrakis A, Plasterk RH, Sixma TK. Crystal structure of the specific DNA-binding domain of Tc3 transposase of C.elegans in complex with transposon DNA. Embo j. 1997 Oct 1;16(19):6044–54.

60. Jung C, Bandilla P, von Reutern M, Schnepf M, Rieder S, Unnerstall U, et al. True equilibrium measurement of transcription factor-DNA binding affinities using automated polarization microscopy. Nat Commun. 2018 Apr 23;9(1):1605.

61. Etheve L, Martin J, Lavery R. Protein-DNA interfaces: a molecular dynamics analysis of time-dependent recognition processes for three transcription factors. Nucleic Acids Res. 2016 Nov 16;44(20):9990–10002.

62. Yesudhas D, Anwar MA, Panneerselvam S, Kim HK, Choi S. Evaluation of Sox2 binding affinities for distinct DNA patterns using steered molecular dynamics simulation. FEBS Open Bio. 2017 Nov;7(11):1750–67.

63. Arnott S. The geometry of nucleic acids. Prog Biophys Mol Biol. 1970;21:265–319.

64. Stofer E, Lavery R. Measuring the geometry of DNA grooves. Biopolymers. 1994 Mar;34(3):337–46.

65. Lide DR. CRC Handbook of Chemistry and Physics, 89th Edition [Internet]. Taylor & Francis; 2008. Available from: https://books.google.com.br/books?id=KACWPwAACAAJ

66. Schulze S, Adams Z, Cerletti M, De Castro R, Ferreira-Cerca S, Fufezan C, et al. The Archaeal Proteome Project advances knowledge about archaeal cell biology through comprehensive proteomics. Nature Communications. 2020 Jun 19;11(1):3145.

67. Bischerour J, Chalmers R. Base flipping in tn10 transposition: an active flip and capture mechanism. PLoS One. 2009 Jul 10;4(7):e6201.

68. van den Broek B, Lomholt MA, Kalisch SMJ, Metzler R, Wuite GJL. How DNA coiling enhances target localization by proteins. Proc Natl Acad Sci U S A. 2008 Oct 14;105(41):15738–42.

69. Cheng X, Roberts RJ. AdoMet-dependent methylation, DNA methyltransferases and base flipping. Nucleic Acids Res. 2001 Sep 15;29(18):3784–95.

70. Fogg JM, Judge AK, Stricker E, Chan HL, Zechiedrich L. Supercoiling and looping promote DNA base accessibility and coordination among distant sites. Nat Commun. 2021 Sep 28;12(1):5683.

71. Priyakumar UD, MacKerell AD. Base Flipping in a GCGC Containing DNA Dodecamer: A Comparative Study of the Performance of the Nucleic Acid Force Fields, CHARMM, AMBER, and BMS. J Chem Theory Comput. 2006 Jan;2(1):187– 200.

72. Randall GL, Zechiedrich L, Pettitt BM. In the absence of writhe, DNA relieves torsional stress with localized, sequence-dependent structural failure to preserve B-form. Nucleic Acids Res. 2009 Sep;37(16):5568–77.

73. Laughton CA, Harris SA. The atomistic simulation of DNA. WIREs Computational Molecular Science. 2011 Jul 1;1(4):590–600.

74. Richardson JM, Dawson A, O’Hagan N, Taylor P, Finnegan DJ, Walkinshaw MD. Mechanism of Mos1 transposition: insights from structural analysis. Embo j. 2006 Mar 22;25(6):1324–34.

75. Reymer A, Zakrzewska K, Lavery R. Sequence-dependent response of DNA to torsional stress: a potential biological regulation mechanism. Nucleic Acids Res. 2018 Feb 28;46(4):1684–94.

76. Teves SS, Henikoff S. DNA torsion as a feedback mediator of transcription and chromatin dynamics. Nucleus. 2014 May;5(3):211–8.

77. Lié O, Renault S, Augé-Gouillou C. SETMAR, a case of primate co-opted genes: towards new perspectives. Mob DNA. 2022 Apr 8;13(1):9.

78. Tellier M, Chalmers R. Human SETMAR is a DNA sequence-specific histone-methylase with a broad effect on the transcriptome. Nucleic Acids Res. 2019 Jan 10;47(1):122–33.

